# GESIAP: A Versatile Genetically Encoded Sensor-based Image Analysis Program

**DOI:** 10.1101/2022.10.05.511006

**Authors:** W. Sharon Zheng, Yajun Zhang, Roger E. Zhu, Peng Zhang, Smriti Gupta, Limeng Huang, Deepika Sahoo, Kaiming Guo, Matthew E. Glover, Krishna C. Vadodaria, Mengyao Li, Tongrui Qian, Miao Jing, Jiesi Feng, Jinxia Wan, Philip M. Borden, Farhan Ali, Alex C. Kwan, Li Gan, Li Lin, Fred H. Gage, B. Jill Venton, Jonathan S. Marvin, Kaspar Podgorski, Sarah M. Clinton, Miaomiao Zhang, Loren L. Looger, Yulong Li, J. Julius Zhu

**Affiliations:** Department of Pharmacology, University of Virginia School of Medicine, Charlottesville, VA 22908; Department of Biomedical Engineering Class 2021, University of Virginia School of Medicine, Charlottesville, VA 22908; Math, Engineering & Science Academy Class of 2025, Albemarle High School, Charlottesville, VA 22901; Biomedical Engineering Class 2024, University of Virginia School of Medicine, Charlottesville, VA 22908; School of Pharmaceutical Sciences, Wenzhou Medical University, Wenzhou 325035, China; School of Neuroscience, Virginia Polytechnic Institute and State University, Blacksburg, VA 24061; Laboratory of Genetics, Salk Institute for Biological Studies, La Jolla, CA 92037; State Key Laboratory of Membrane Biology and Peking-Tsinghua Center for Life Sciences, Peking University, Beijing 100871, China; Janelia Research Campus, Howard Hughes Medical Institute, Ashburn, VA 20147; Department of Psychiatry, Yale University School of Medicine, New Haven, CT 06511; Helen and Robert Appel Alzheimer’s Disease Research Institute, Weill Cornell Medicine College, New York, NY 10065; Department of Chemistry, University of Virginia, Charlottesville, VA 22908; Department of Electrical and Computer Engineering, University of Virginia, Charlottesville, VA 22908

**Keywords:** Algorithms, genetically encoded fluorescent sensor, neuromodulation, quantal analysis, synaptic transmission

## Abstract

Intercellular communication mediated by a large number of neuromodulators diversifies physiological actions, yet neuromodulation remains poorly understood despite the recent upsurge of genetically encoded transmitter sensors. Here, we report the development of a versatile genetically encoded sensor-based image analysis program (GESIAP) that utilizes MATLAB-based algorithms to achieve high-throughput, high-resolution processing of sensor-based functional imaging data. GESIAP enables delineation of fundamental properties (e.g., transmitter spatial diffusion extent, quantal size, quantal content, release probability, pool size, and refilling rate at single release sites) of transmission mediated by various transmitters (i.e., monoamines, acetylcholine, neuropeptides, and glutamate) at various cell types (i.e., neurons, astrocytes, and other non-neuronal cells) of various animal species (i.e., mouse, rat, and human). Our analysis appraises a dozen of newly developed transmitter sensors, validates a conserved model of restricted non-volume neuromodulatory synaptic transmission, and accentuates a broad spectrum of presynaptic release properties that variegate neuromodulation.

## INTRODUCTION

Intercellular communication mediated by fast-acting transmitters glutamate and gamma-aminobutyric acid (**GABA**), and by hundreds of slow-acting neuromodulatory transmitters (e.g., monoamines, neuropeptides, and other molecules), orchestrates diverse behavioral and physiological phenomena (Greengard, 2001; Sudhof, 2018). Sustained interest in understanding intercellular communication stems from the belief that this knowledge will not only advance our understanding of the brain and body, but also unveil pathogenic mechanisms of and therapeutic strategies for diverse neurological and psychiatric disorders, as well as other health conditions. Indeed, structural and functional alterations in fast transmission are commonly seen in many symptomatically distinct disorders (Bayes et al., 2011; Volk et al., 2015; Wichmann and Kuner, 2022), and aberrant synaptic communication have been identified as the underlying cause of a number of diseases (Henley and Wilkinson, 2016; Sudhof, 2018). Understanding of the mechanisms of fast transmission has been gained through the remarkable sensitivity and temporal resolution of electrophysiological recordings and herculean experiments that have resolved every synaptic property of glutamatergic and GABAergic transmission (von Gersdorff and Borst, 2002; Wu et al., 2014; Pulido and Marty, 2017). These properties define transmission, including its precision (e.g., spatial diffusion extent), strength (e.g., quantal size and quantal content), sustainability (e.g., vesicular refilling rate), and short- and long-term plasticity (e.g., release probability) of intercellular communication (Kaeser and Regehr, 2014; Jackman and Regehr, 2017). Unfortunately, patch-clamp recordings are compromised when examining cells with minimal and/or desensitizing neuromodulator-induced electrophysiological responses (Dani and Bertrand, 2007; Nadim and Bucher, 2014). Therefore, despite years of research that implicates numerous slow-acting neuromodulatory transmitters in myriad physiological actions and diseases, there are few quantitative and mechanistic insights into neuromodulatory regulation and function.

Recently developed genetically encoded sensors for neuromodulatory transmitters provide new opportunities to monitor and quantitate neuromodulation (Sabatini and Tian, 2020; Labouesse and Patriarchi, 2021; Lin et al., 2021; Wu et al., 2022). Indeed, both theoretical (Lin et al., 2021) and preliminary experimental (Zhu et al., 2020) analyses have shown that several genetically encoded transmitter sensors emit a large number of photons following transmitter binding to enable high-resolution visualization of the process. However, most genetically encoded transmitter sensors remain underused, due mainly to the lack of user-friendly analysis programs for high-sensitivity, high-resolution visualization of transmitter release. Here, we report the development of a genetically encoded sensor-based image analysis program (**GESIAP**) that optimizes and integrates MATLAB-based alignment, deconvolution, baseline correction, denoise, and background subtraction algorithms into a single automatic analysis program to maximize its image analysis sensitivity, resolution, and efficiency. GESIAP enables high-sensitivity and high-resolution visualization of transmission mediated by various transmitters (i.e., monoamines, acetylcholine, neuropeptides, and glutamate) at various cell types (i.e., neurons, astrocytes, and other non-neuronal cells) of various animal species (i.e., mouse, rat, and human). The GESIAP-enabled visualization of transmitter release immediately allows decoding of fundamental synaptic properties, including transmitter spatial diffusion extent, quantal size, quantal content, release probability, pool size, and refilling rate at single release sites with a few simple sets of imaging experiments, much more efficiently than patch-clamp recordings. Our data illustrate a functional scheme of neuromodulatory transmitters that share a conserved, restricted non-volume transmission mode and yet, differ extensively in presynaptic release properties that could diversify neuromodulatory actions. GESIAP is poised to be the first versatile tool applicable in hypothesis-based experiments that can causally link neuromodulatory transmission properties with physiological actions and clinical symptoms in diseases.

## RESULTS

### Development of GESIAP

To study properties of intercellular neuromodulatory communication, we developed GESIAP to enhance visualization of transmitter release, taking advantage of the high signal-to-noise ratio of modern genetically encoded transmitter sensors (Lin et al., 2021) and expeditiously improved computational algorithms (Arigovindan et al., 2013; Weigert et al., 2018; Koho et al., 2019). GESIAP consists of five major algorithmic procedures to perform alignment, deconvolution, baseline correction, denoise, and background subtraction for fluorescence image sequences collected from genetically encoded sensor-based functional imaging experiments (**Fig 1A-B**). After several dozen rounds of iterative improvements in the major algorithmic procedures, GESIAP can now achieve nanoscopic visualization of transmitter release monitored with genetically encoded transmitter sensors (**Fig 1A-B**). GESIAP is deposited at UVA Patent Foundation (https://lvg.virginia.edu/Zhu_lab/GESIAP/) (currently password protected) and will be freely available for general users in the research community.

**Figure 1.**
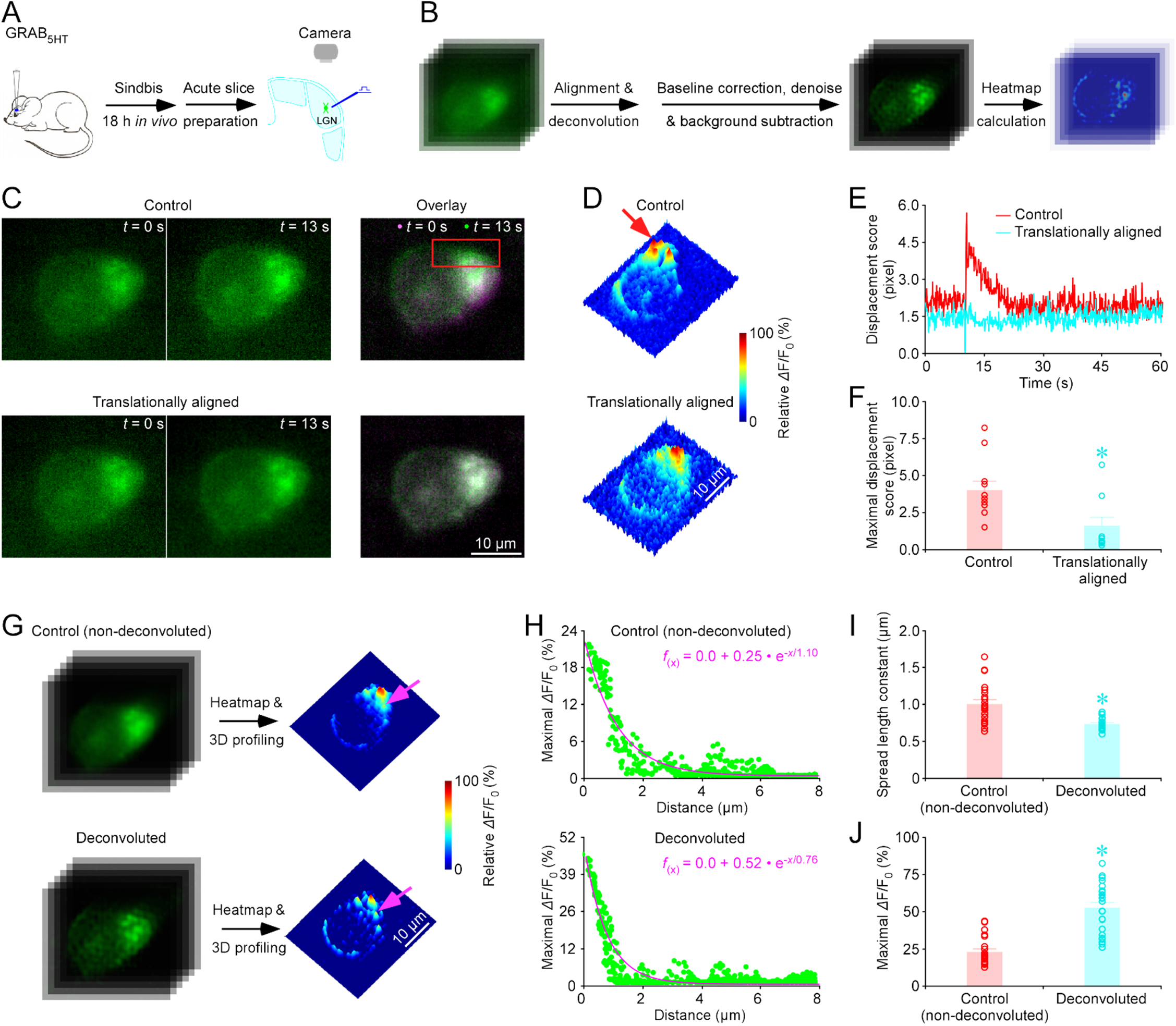
Translational alignment and deconvolution improve image faithfulness and resolution. (**A**) Schematic of stimulation-imaging experiment in an *ex vivo* mouse geniculate preparation. LGN: the lateral geniculate nucleus. (**B**) GESIAP integrates the procedures performing translational alignment, Landweber deconvolution, baseline correction, denoise, background subtraction, *Δ*F/F_0_ heatmap calculation, and three-dimensional (**3D**) transmitter release profiling. (**C**) Snapshots and overlay of images captured at two time points of a GRAB_5HT1.0_ expressing geniculate neuron. Note the prominent mismatch in the overlay of control unaligned images. (**D**) 3D spatiotemporal profiling of electrically evoked fluorescence *Δ*F/F_0_ responses in the GRAB_5HT1.0_ expressing geniculate neuron shown in **C**. Note the spurious peaks from motion artifacts (red arrow indicates one such peak). (**E**) Displacement score of the geniculate neuron. (**F**) Maximal displacement scores before and after translational alignment (Before: 4.2±0.7 vs. After: 1.7±0.6 *n* = 10 neurons, *Z* = −2.803 *p* = 0.002). (**G**) Snapshots (left) and the corresponding 3D spatiotemporal profiling (right) of electrically evoked fluorescence *Δ*F/F_0_ responses in the GRAB_5HT1.0_ expressing geniculate neuron under control (non-deconvolved) and deconvolved conditions. (**H**) Pixel-wise maximal *Δ*F/F_0_ at the isolated release site indicated by the pink arrow in **G**. Fitting the data points with a single-exponential decay function (pink line) yields an estimated serotonin spatial spread length constant of 1.10 μm for non-deconvolved images and 0.76 μm for deconvolved images. (**I**) Average spatial spread length constants for serotonin at geniculate neurons obtained from control nondeconvolved and deconvolved images (Control: 1.01±0.06 μm vs. Deconvolved: 0.74±0.02 μm; *n* = 22 from 8 neurons, *Z* = −4.042, *p* < 0.001). (**J**) The maximum *Δ*F/F_0_ responses obtained from control non-deconvolved and deconvolved image (Control: 23.2±2.0% vs. Deconvolved: 52.4±3.5%; *n* = 22 from 8 neurons, *Z* = 4.107, *p* < 0.001). Asterisks indicate *p* < 0.05 (Wilcoxon tests).

### GESIAP enables visualization of synaptic transmission

We initially tested the applicability of GESIAP using an *ex vivo* brain slice preparation with Sindbis viral expression of a genetically encoded sensor for serotonin, GRAB_5HT1.0_ (Wan et al., 2021), in thalamic geniculate neurons in intact mice. Then we prepared acute thalamic slices after ~18 hours of *in vivo* expression (**Fig 1A**). In the acute thalamic slices, we activated serotonergic fibers with local electric stimulation to evoke fluorescence responses at GRAB_5HT1.0_ expressing geniculate neurons and then used GESIAP to visualize serotonin release (**Fig 1A-B**).

The first GESIAP procedure, image alignment, aims to preserve the veracity of fluorescence responses by correcting drifts, fluctuations, and motion artifacts occurring during microscopic imaging. The iterative translational alignment procedure corrected movements in fluorescence responses (**Fig 1C**), thereby preventing spurious introduction of serotonin release “peaks” caused by movement-induced pixel displacements (**Fig 1D**). Quantitative analysis revealed that translational alignment largely eliminated movement-induced pixel displacements and minimized movement displacement scores (**Fig 1E-F**). Likewise, an affine alignment procedure made similar movement corrections and prevented creation of spurious serotonin release peaks, albeit requiring ~10x longer compute times (**Fig S1**). These results underscore the importance of image alignment in preserving the reliability of fluorescence responses.

The second GESIAP procedure, deconvolution, improves the quality of microscopic images by “reassigning” detected photons to their original emitting locations. GESAIP utilizes Landweber deconvolution procedure (Sage et al., 2017), together with the point spread function empirically measured from our wide-field microscopic system (**Fig S2A-C**), to reverse the optical distortion induced by light diffraction (**Fig 1G-H**). The procedure sharpened the image of individually isolated serotonin release events, and fitting with a single-exponential decay function on pixel-wise maximal *Δ*F/F_0_ plots at single isolated release sites produced serotonin spatial spread length constants (**Fig 1I**). Landweber deconvolution increased the maximal *Δ*F/F_0_ at isolated release sites by over twofold and improved the spatial optical resolution by more than 25% (**Fig 1J-K**). Re-convolving deconvolved images largely reinstated the original microscopic images (**Fig S2D-F**), verifying the quality of deconvolution (cf. (Thomas and Agard, 1984)). We found that ~25-50 iterations of deconvolution essentially maximized image improvement, as indicated by the diminished gain in spread length measurement and minimized difference between original and convolved images (**Fig S2G**). The procedure comes with some tradeoffs in signal-to-noise ratio in the deconvolved images (**Fig S2H**), which can be largely mitigated by denoise (see below).

The subsequent GESIAP algorithms, baseline correction, denoise, and background subtraction, are intended to refine the quality of fluorescence *Δ*F/F_0_ responses (**Fig 2**). In particular, baseline correction largely compensates for the slow reduction in light intensity caused by sensor photobleaching during prolonged imaging processes (**Fig 2A-C**). Denoise essentially eliminates random noise, and background subtraction improves the visibility of *Δ*F/F_0_ responses (**Fig 2A-B and D-F**). Importantly, while these three procedures improved visualization of the *Δ*F/F_0_ responses, particularly responses at individual release sites, none of them affected serotonin spread length constants. Furthermore, background subtraction enhanced maximal *Δ*F/F_0_ by another three-fold at GRAB_5HT1.0_ expressing geniculate neurons (**Fig 2G-H**). Together, these results indicate that GESIAP enables high-resolution visualization of evoked serotonergic fluorescence responses at GRAB_5HT1.0_ expressing geniculate neurons.

**Figure 2.**
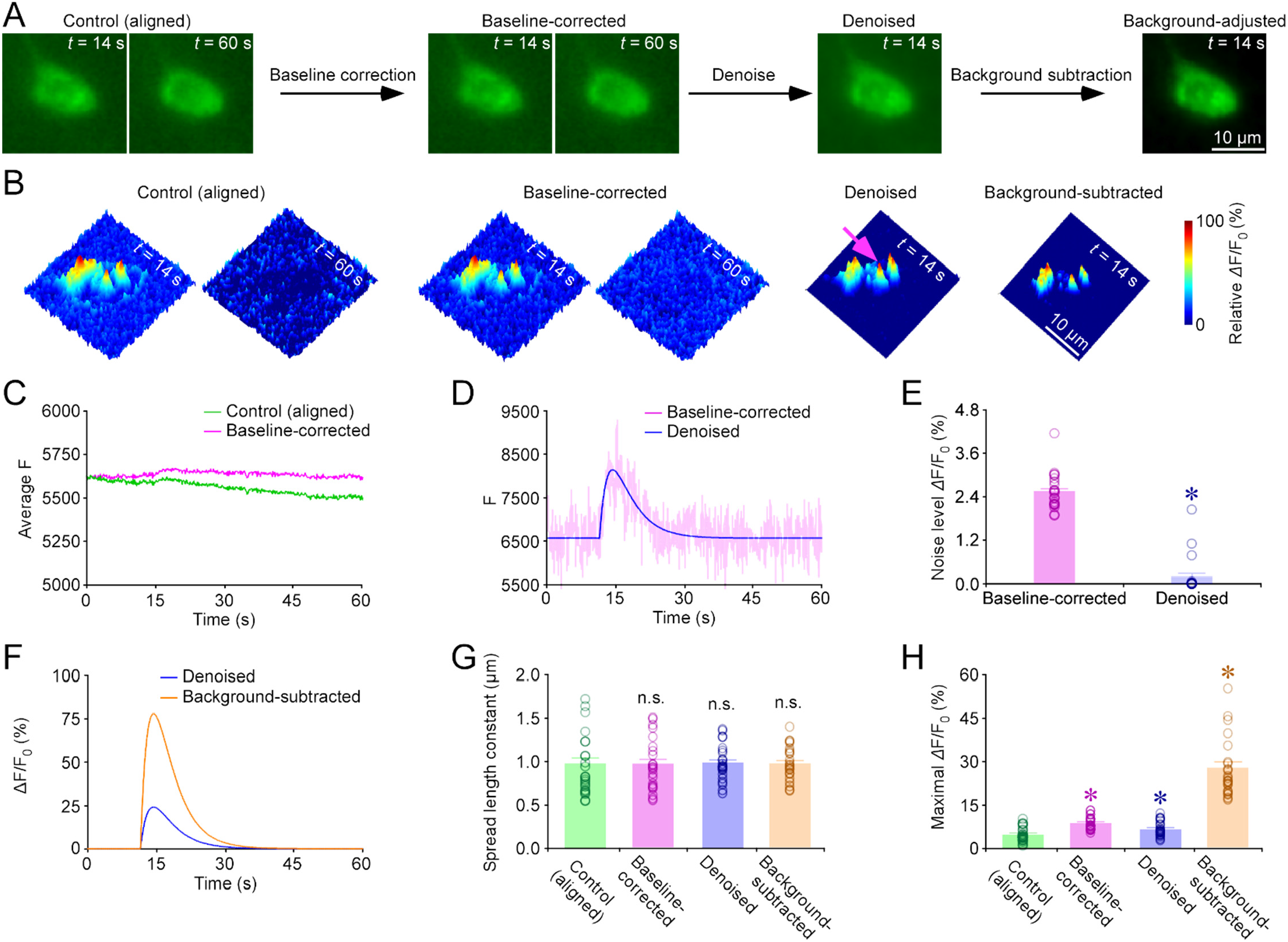
Baseline correction, denoise, and background subtraction refine image quality and appearance. (**A-B**) Snapshots (**A**) and 3D spatiotemporal profiling (**B**) of electrically evoked fluorescence *Δ*F/F_0_ responses in a GRAB_5HT1.0_ expressing geniculate neuron under control (aligned), baseline-corrected, denoised, and background-subtracted conditions. (**C**) Plot of average fluorescence responses of the cell over time for control baseline-uncorrected and baseline-corrected images. Note the minimal photobleaching in the baseline-corrected images. (**D**) Plot of fluorescence responses at the point indicated by the pink arrow in **B** over time with baseline-corrected and further denoised with double-exponential synaptic function. (**E**) Noise level measured at the peaks of control baseline-corrected and further denoised images (Control: 2.8±0.1% vs. Denoised: 0.2±0.1%; *n* = 22 from 8 neurons, *Z* = −4.107, *p* < 0.001). (**F**) Plot of *Δ*F/F_0_ responses at the point indicated by the pink arrow in **B** over time with baseline-corrected and further denoised with double-exponential synaptic function. (**G**) Average spatial spread length constants for serotonin at geniculate neurons obtained from baseline-corrected, denoised, and background-adjusted images (Baseline-corrected: 0.97±0.06 μm, *Z* = 0.406, *p* = 0.679; denoised: 0.99±0.04 μm, *Z* = 1.023, *p* = 0.314; background-subtracted: 0.98±0.04 μm, *Z* = 828, *p* = 0.417) compared to that of control (aligned) (Control: 0.96±0.07 μm; *n* = 22 from 8 neurons). (**H**) Maximal *Δ*F/F_0_ responses obtained from baseline-corrected, denoised, and background-subtracted images (Baseline-corrected: 8.9±0.5%, *Z* = 2.906, *p* = 0.004; denoised: 6.7±0.3%, *Z* = −3.393, *p* < 0.001; background-subtracted: 28±2%, *Z* = 4.107, *p* < 0.001) compared to control (aligned) (Control: 8.3±0.5%; *n* = 22 from 8 neurons). Asterisks indicate *p* < 0.05 (Wilcoxon tests).

### GESIAP permits resolution of fundamental synaptic properties

Visualization of fluorescence responses at isolated transmitter release sites permits decoding of fundamental properties of synaptic transmission (**Fig 3** and **Movie S1A-B**). GESIAP procedures revealed that, on average, local electric stimulation evoked serotonin release at ~10 sites on somata of geniculate neurons (**Fig 3A-D**). We analyzed the spatial diffusion of serotonin at those well-isolated release sites (**Fig 3C**). Fitting with a single-exponential decay function on pixel-wise maximal *Δ*F/F_0_ plots at isolated serotonin release sites yielded a serotonin spread length constant of ~0.75 μm at geniculate neurons (**Fig 3E-F**). To control for potential expression-level effects, analysis showed a weak correlation between *Δ*F/F_0_ responses and basal fluorescence F_0_ (see **Methods;** cf. (Zhu et al., 2020)), suggesting that *Δ*F/F_0_ responses were largely independent of GRAB_5HT1.0_ expression levels. Bath application of tetrodotoxin (**TTX**), which blocks action potential-dependent synaptic transmission, eliminated the evoked responses at GRAB_5HT1.0_ expressing geniculate neurons (**Fig S3**), confirming the synaptic origin of signals (cf. (Zhu et al., 2020; Wan et al., 2021)). These results suggest that the evoked serotonin release is spatially restricted, non-volume synaptic transmission.

**Figure 3.**
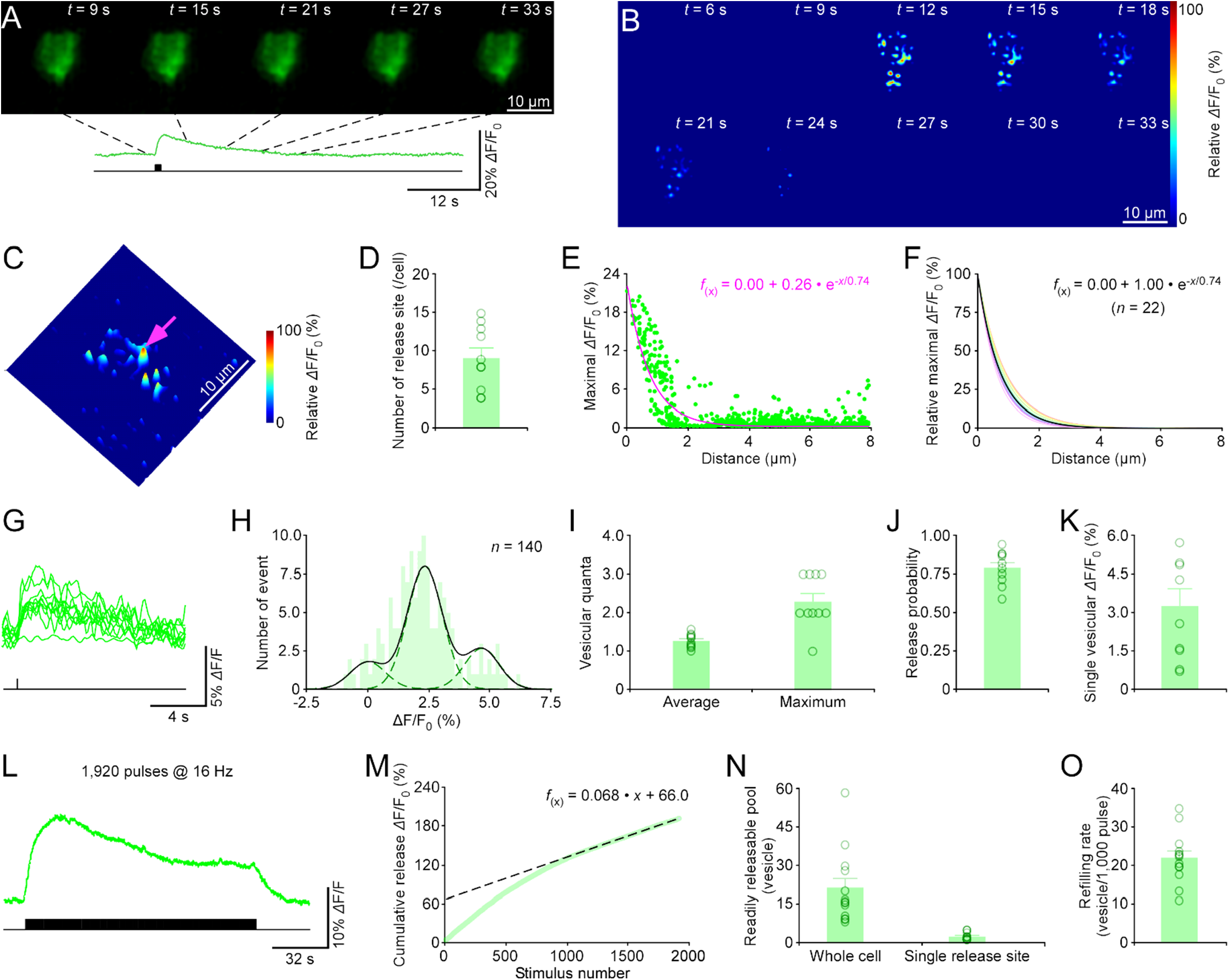
GESIAP enables resolution of serotonergic synaptic properties at geniculate neurons. (**A-C**) Snapshots (**A**), heatmaps (**B**), and 3D spatiotemporal profiling (**C**) of electrically evoked fluorescence responses in a GRAB_5HT1.0_ expressing geniculate neuron. Note one isolated release site indicated by a pink arrow in **C**. (**D**) Average number of release sites of geniculate neurons (9.2±1.3; *n* = 8 neurons). (**E**) Pixel-wise maximal *Δ*F/F_0_ at the isolated release site indicated by pink arrow in **C**. Fitting the data points in this plot with a single-exponential decay function (pink line) yields an estimated serotonin spatial spread length constant of 0.74 μm. (**F**) Summary plot of 5HT diffusion curves from putative single release sites gives an average spatial spread length constant of 0.74±0.02 μm for serotonin at geniculate neurons (*n* = 22 from 8 neurons). Note the average diffusion curve in black. (**G**) Ten *Δ*F/F_0_ responses evoked by single pulse stimuli at the isolated release site (indicated by the pink arrow in **C**) of the GRAB_5HT1.0_ expressing geniculate neuron. (**H**) Amplitude histograms of *Δ*F/F_0_ responses show multiple, nearly equally spaced peaks. (**I-K**) Values for the average (1.26±0.06) and maximal (2.10±0.18) vesicular quanta, release probability (0.79±0.04), and quantal size (3.2±0.6 %*Δ*F/F_0_) at single serotonin release sites. **(L**) Fluorescence *Δ*F/F_0_ responses of a GRAB_5HT1.0_ expressing geniculate neuron induced by local electrical stimuli of 1,920 pulses at 16 Hz. **(M**) Plots of cumulative fluorescence *Δ*F/F_0_ responses against stimulus number in the GRAB_5HT1.0_ expressing geniculate neuron. Note the readily releasable pool size and refilling rate inferred from the *y*-axis intercept and slope of a linear fit to late points of the cumulative trace, respectively. (**N-O**) Values for the readily releasable pool (20.8±4.0, *n* = 13 neurons for whole cells; 2.7±0.4; *n* = 13 neurons for single release sites) and vesicular refilling rate (2.2±0.2 vesicles/per 100 pulses; *n* =13 neurons) of GRAB_5HT1.0_ expressing geniculate neurons.

The ability to visualize serotonin release at isolated subcellular locations with the nanoscale resolution allows decoding of presynaptic properties of serotonergic transmission (**Fig 3G-K** and **Movie S1C**). We first examined *Δ*F/F_0_ responses at single release sites of GRAB_5HT1.0_ expressing geniculate neurons evoked by locally delivered trains of electric pulses at 0.1 Hz. The evoked responses at single releases sites appeared as stochastic events, exhibiting failures and releases of *Δ*F/F_0_ events of varied sizes (**Fig 3G-H**). Plotting a *Δ*F/F_0_ amplitude histogram, an analysis method adapted from the classic quantal analysis approach (Fatt and Katz, 1952), revealed multiple, nearly equally spaced peaks, revealing the number of released vesicles (average ~1.25 vesicular quanta and up to 3 vesicular quanta per stimulus) (**Fig 3G-I**). In addition, the analysis enabled direct readout of a release probability *Pr* of ~0.75 (release success rate over multiple trials) at single release sites and a quantal size of ~3% *Δ*F/F_0_ for serotonergic vesicles (**Fig 3H-K**). We then applied a prolonged stimulus consisting of 1,920 pulses at GRAB_5HT1.0_ expressing geniculate neurons, with the initial pulses evoking larger *Δ*F/F_0_ responses followed by plateaued *Δ*F/F_0_ responses at later pulses (**Fig 3L**). Plotting the cumulative release against stimulus number and fitting the plot with a previously established synaptic transmission model (Neher, 2015) revealed multiple synaptic properties of serotonergic termini (**Fig 3M**). In particular, readily releasable pool size and refilling rate were inferred from back-extrapolation and slope of a linear fit to late points of the cumulative trace, respectively (Schneggenburger et al., 1999). Converting the *Δ*F/F_0_ values to quantal size (**Fig 3K**) resulted in whole cell and single release pool sizes of ~55 and ~5 vesicles, respectively, and a refilling rate of ~20 vesicles per 1,000 pulses (**Fig 3N-O**). Collectively, nanoscale visualization of serotonin release permits resolution of the number of serotonin release sites and spatial diffusion extent, quantal size, quantal content, release probability, refilling rate and vesicular pool size at GRAB_5HT1.0_ expressing geniculate neurons.

### GESIAP enables visualization of transmission of various animal species

We wished to validate the general applicability of GESIAP across preparations. Hence, we imaged serotonergic transmission in another mouse brain area, the dorsal raphe nucleus by mediating Sindbis viral expression of GRAB_5HT1.0_ in the nucleus *in vivo* and preparing acute raphe slices after ~18 hours of expression (**Fig S4A**). In acute raphe slices, local electric stimulation evoked fluorescence *Δ*F/F_0_ responses at GRAB_5HT1.0_ expressing raphe neurons, and GESIAP enabled visualization of *Δ*F/F_0_ responses (**Fig S4B-D**). Analysis of the spatial diffusion extent of serotonin yielded a serotonin spread length constant of ~0.70 μm at mouse raphe neurons (**Fig S4E-F**). To test the applicability of GESIAP in another animal species, we repeated the same experiment using an *ex vivo* rat hippocampal preparation to visualize the evoked serotonergic fluorescence responses at dentate gyrus neurons (**Fig S5A-D**). GESIAP produced a serotonin spread length constant of ~0.75 μm at dentate gyrus neurons of rats (**Fig S5E-F**). Together, these data suggest that GESIAP is applicable for visualization of serotonergic transmission at rodent neurons in general.

Serotonin is involved in a wide array of human behaviors and diseases (Okaty et al., 2019). We investigated human serotonergic transmission using an established human fibroblast-derived serotonergic neuron preparation (Vadodaria et al., 2016) (**Fig 4A-B**). Local electric stimulation evoked fluorescence responses at GRAB_5HT1.0_ expressing human neurons, and GESIAP visualized the responses and provided a serotonin spread length constant of ~0.70 μm at human fibroblast-derived neurons (**Fig 4C-G**). Collectively, these results indicate that GESIAP enables visualization of evoked serotonergic fluorescence *Δ*F/F_0_ responses at distinct types of neurons of different animal species, all of which display restricted, non-volume synaptic transmission.

**Figure 4.**
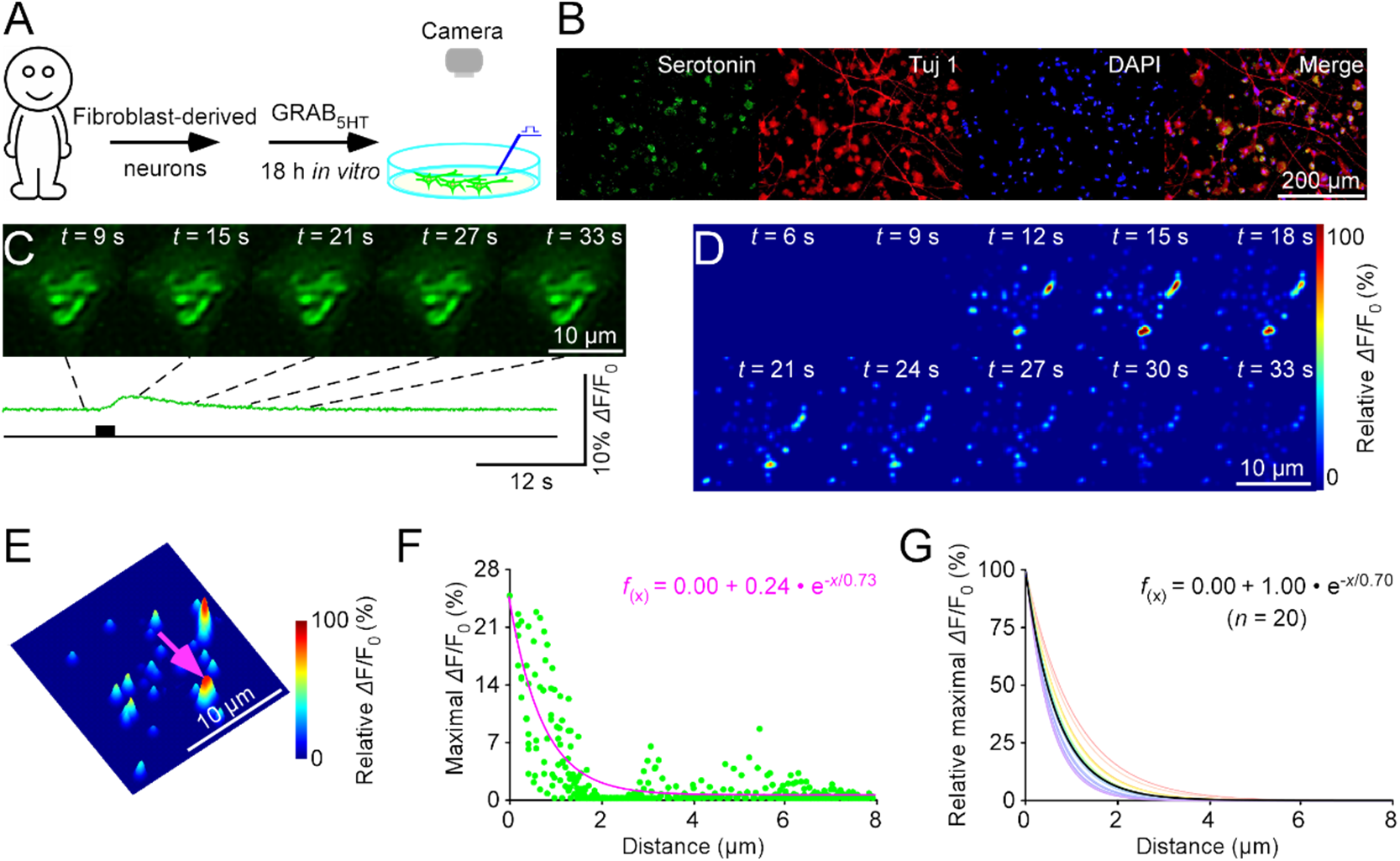
Nanoscopic visualization of serotonergic transmission at human neurons. (**A**) Schematic of stimulation-imaging experiment in a human fibroblast-derived neuron culture preparation. (**B**) Immunostaining of human fibroblast-derived serotoninergic neurons. Serotonin: anti-serotonin staining; Tuj1: anti-tubulin β3 staining; DAPI: 4’,6-diamidino-2-phenylindole nucleic acid staining. (**C-E**) Snapshots (**C**), heatmaps (**D**), and 3D spatiotemporal profiling (**E**) of electrically evoked *Δ*F/F_0_ responses in a GRAB_5HT1.0_ expressing human fibroblast-derived neuron. Note the isolated release site indicated by a pink arrow in **E**. (**F**) Pixel-wise maximal *Δ*F/F_0_ at the isolated release site indicated by the pink arrow in **E**. Fitting the data points with a single-exponential decay function (pink line) yields an estimated serotonin spatial spread length constant of 0.73 μm. (**G**) Summary plot of serotonin diffusion curves from putative single release sites gives an average spatial spread length constant of 0.70±0.03 μm for serotonin at human fibroblast-derived neurons (*n* = 20 from 11 neurons). Note the average diffusion curve in black.

### GESIAP enables visualization of transmission at both neuronal and non-neuronal cells

We next wanted to learn whether GESIAP applicability could be extended beyond neuronal cells. Cholinergic transmission regulates physiology of many neuronal and non-neuronal cells (Picciotto et al., 2012; Ballinger et al., 2016; Herring et al., 2019). We validated applicability in analyzing cholinergic transmission by expressing a genetically encoded acetylcholine sensor, iAChSnFR (Borden et al., 2020; Zhu et al., 2020), in layer 2 (**L2**) of the intact mouse medial entorhinal cortex using Sindbis (**Fig S6A**). Approximately 18 hours after *in vivo* Sindbis viral expression, we made acute entorhinal slices and imaged fluorescence responses of iAChSnFR expressing entorhinal L2 stellate neurons to electrical stimulation of entorhinal cortical L1, a layer densely innervated by cholinergic fibers originating from the basal forebrain (Ray et al., 2014) (**Fig S6B**). The evoked cholinergic responses were processed by GESIAP, which provided high-resolution visualization of cholinergic transmission and resolved an acetylcholine spread length constant of ~0.70 μm at entorhinal L2 stellate neurons (**Fig S6C-G**). Next, we investigated cholinergic transmission at astrocytes, which express high levels of muscarinic ACh receptors in their fine distal processes involved in tripartite synapses (Van Der Zee et al., 1993). We used a lentiviral vector carrying a glial fibrillary acidic protein (**GFAP**) promoter to achieve targeted expression of iAChSnFR in entorhinal astrocytes in mice *in vivo* for seven days and imaged an electrically-evoked *Δ*F/F_0_ responses *ex vivo* in acute entorhinal brain slices (**Fig 5A-B**). Similarly, GESIAP allowed high-resolution visualization of cholinergic transmission, and resolved an acetylcholine spread length constant of ~0.70 μm at entorhinal L2 astrocytes (**Fig 5C-G**). These results suggest that GESIAP allows visualization of cholinergic transmission at both neurons and non-neuronal cells in the brain.

**Figure 5.**
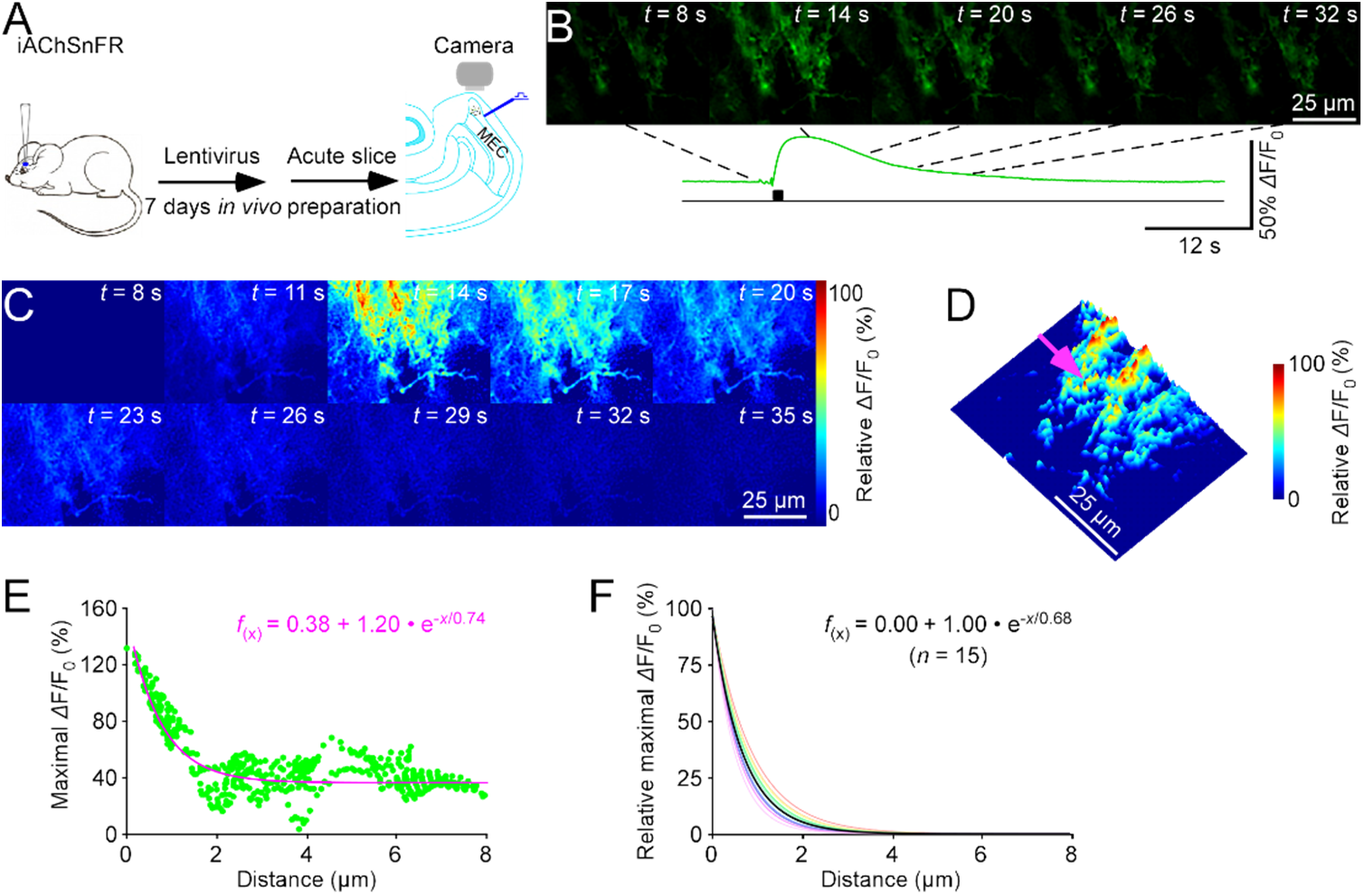
Nanoscopic visualization of cholinergic transmission at entorhinal astrocytes. (**A**) Schematic of stimulation-imaging experiment in an *ex vivo* entorhinal preparation. MEC: the medial entorhinal cortex. (**B-D**) Snapshots (**B**), heatmaps (**C**), and 3D spatiotemporal profiling (**D**) of electrically evoked *Δ*F/F_0_ responses in an iAChSnFR expressing entorhinal astrocyte. Note the isolated release site indicated by a pink arrow in **D**. (**E**) Pixel-wise maximal *Δ*F/F_0_ at the isolated release site indicated by the pink arrow in **D**. Fitting the data points with a single-exponential decay function (pink line) yields an estimated acetylcholine spatial spread length constant of 0.74 μm. (**F**) Summary plot of acetylcholine diffusion curves from putative single release sites gives an average spatial spread length constant of 0.70±0.02 μm for acetylcholine at entorhinal astrocytes (*n* = 22 from 12 cells). Note the average diffusion curve in black.

We then investigated cholinergic release in the mouse pancreas and adrenal gland (**Fig S7A and S8A**), in which parasympathetically released acetylcholine controls insulin secretion and regulates blood pressure and steroid release, respectively (Ungar and Phillips, 1983; Satin and Kinard, 1998). We introduced another acetylcholine sensor, GRAB_ACh2.0_ (Jing et al., 2018), in the pancreas and adrenal gland *in vivo* with Sindbis, and we imaged fluorescence responses of GRAB_ACh2.0_ expressing cells in acute pancreatic and adrenal slices after 18 hours expression (**Fig S7A-B and S8A-B**). GESIAP permitted high-resolution visualization of acetylcholine release, and calculated an acetylcholine spread length constant of ~0.75 μm at both pancreatic and adrenal cells (**Fig S7C-G and S8C-G**). Collectively, these results show that GESIAP is applicable for visualization of transmission at both neuronal and non-neuronal cells.

### GESIAP enables visualization of transmission mediated by various transmitters

We next tested whether GESIAP is applicable for visualization of transmission mediated by transmitters other than serotonin and acetylcholine. We began with Sindbis expression of a genetically encoded norepinephrine sensor, GRABNE1m (Feng et al., 2019), in the mouse amygdala (**Fig 6A**), which is heavily innervated by noradrenergic fibers from the locus coeruleus (Foote et al., 1983). About 18 hours later, we prepared acute amygdalar slices and electrically evoked adrenergic fluorescence responses at GRABNE1m expressing neurons (**Fig 6B**). GESIAP visualized norepinephrine release, and calculated a norepinephrine spread length constant of ~0.70 μm at amygdalar neurons (**Fig 6C-G**). Similarly, we employed Sindbis virus for *in vivo* expression of a genetically encoded dopamine sensor, GRAB_DA2.0_, in the mouse striatum (**Fig S9A**). Around 18 hours later, we prepared acute striatal slices and electrically evoked dopaminergic fluorescence responses at GRAB_DA2.0_ expressing neurons (**Fig S9B**). GESIAP visualized dopamine release, and calculated a dopamine spread length constant of ~0.70 μm at striatal neurons (**Fig S9C-G**). Finally, we used GESIAP to visualize histamine release at the amygdala, and calculated a histamine spread length constant of ~0.70 μm at amygdalar neurons (**Fig S10**). These results suggest that GESIAP is applicable for visualization of transmission mediated by acetylcholine and monoamines in general.

**Figure 6.**
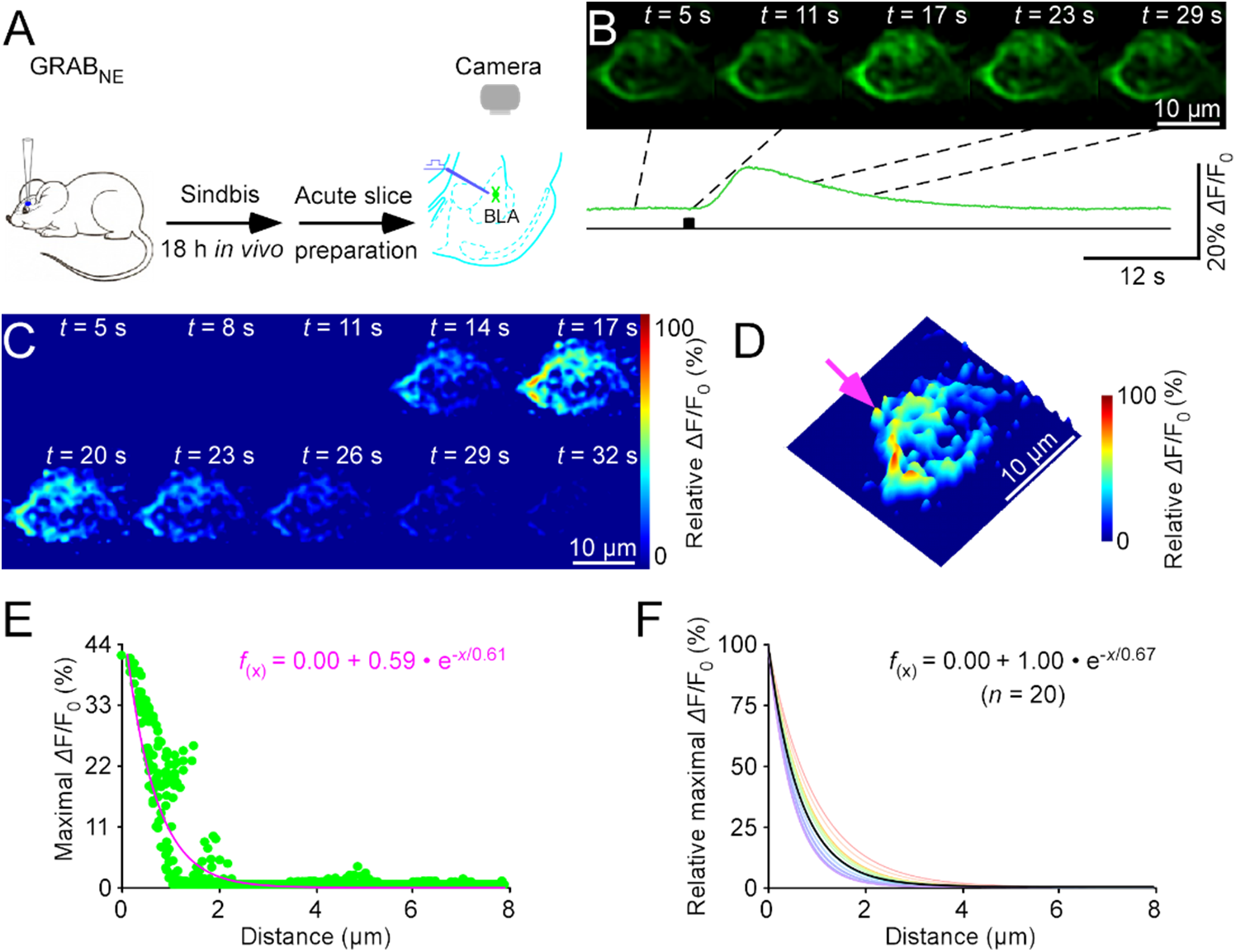
Nanoscopic visualization of adrenergic transmission at amygdalar neurons. (**A**) Schematic of stimulation-imaging experiment in an *ex vivo* amygdalar preparation. BLA: the basolateral amygdala. (**B-D**) Snapshots (**B**), heatmaps (**C**), and 3D spatiotemporal profiling (**D**) of electrically evoked *Δ*F/F_0_ responses in a GRAB_NE1.0_ expressing amygdalar neuron. Note the isolated release site indicated by a pink arrow in **D**. (**E**) Pixel-wise maximal *Δ*F/F_0_ at the isolated release site indicated by the pink arrow in **D**. Fitting the data points with a single-exponential decay function (pink line) yields an estimated norepinephrine spatial spread length constant of 0.61 μm. (**G**) Summary plot of norepinephrine diffusion curves from putative single release sites gives an average spatial spread length constant of 0.67±0.03 μm for norepinephrine at amygdalar neurons (*n* = 20 from 10 neurons). Note the average diffusion curve in black.

Finally, we studied whether GESIAP is applicable for visualization of transmission mediated by neuropeptides, which belong to the largest group of neurotransmitters (>100) and have diverse effects on physiology and behavior (Nusbaum et al., 2017; Guillaumin and Burdakov, 2021). We first expressed a genetically encoded oxytocin sensor, GRAB_OT1.0_ (Qian et al., 2022), in the mouse paraventricular nucleus and ventral tegmental area *in vivo*. Approximately 18 hours later, we prepared acute paraventricular and ventral tegmental slices, and then electrically evoked oxytocin fluorescence responses at expressing neurons (**Fig 7A** and **Movie S2A-B** and **S3A-B**). GESIAP revealed oxytocin release at ~8.5 and ~6.5 sites on somata of paraventricular and ventral tegmental neurons, respectively (**Fig 7B-C**). Calculation of transmitter spatial diffusion extent yielded an oxytocin spread length constant of ~0.75 μm at both paraventricular and ventral tegmental neurons (**Fig 7D-E**). Trains of electric pulses at 0.1 Hz evoked stochastic responses at single releases sites of paraventricular and ventral tegmental neurons (**Fig 7F** and **Movie S2C** and **S3C**). Quantal analysis revealed ~1.25 vesicular quanta and up to 3 vesicular quanta per release at paraventricular neurons, and an average of ~1.75 vesicular quanta and up to 4 vesicular quanta per stimulus at ventral tegmental neurons (**Fig 7G-H**). The analysis also gave a release probability *P*_r_ of ~1.0 at single release sites of both paraventricular and ventral tegmental neurons, and a quantal size of ~12-15% *Δ*F/F_0_ for oxytocinergic vesicles (**Fig 7I-J**). A prolonged stimulus of 1,920 pulses at GRAB_OT1.0_ expressing paraventricular and ventral tegmental neurons largely depleted oxytocin at ventral tegmental neurons, but not at paraventricular neurons (**Fig 7K-L**). Converting *Δ*F/F_0_ responses to quantal size (**Fig 7J**) yielded a whole cell vesicular pool size of ~25 and ~35 vesicles for paraventricular and ventral tegmental neurons, respectively, and a single release site vesicular pool size of ~5 and ~7 vesicles for paraventricular and ventral tegmental neurons, respectively (**Fig 7M**). This yielded a refilling rate of ~20 and ~1 vesicles per 1,000 pulses for paraventricular and ventral tegmental neurons, respectively (**Fig 7N**).

**Figure 7.**
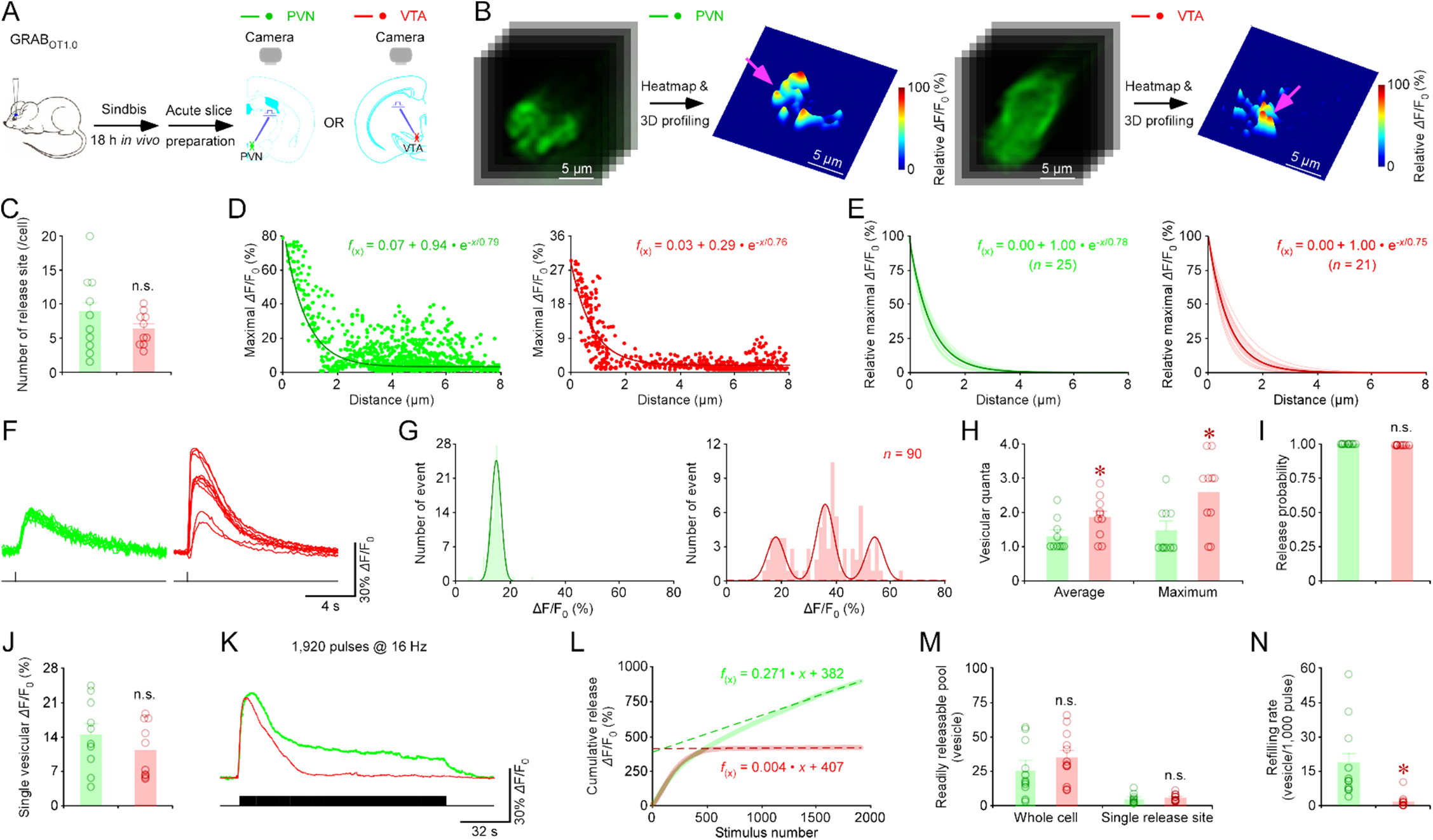
Nanoscopic visualization of oxytocinergic transmission at paraventricular and ventral tegmental neurons. (**A**) Schematic of stimulation-imaging experiment in an *ex vivo* paraventricular and ventral tegmental preparation. PVN: the paraventricular nucleus; VTA: the ventral tegmental area. (**B**) Snapshots and 3D spatiotemporal profiling of electrically evoked fluorescence *Δ*F/F_0_ responses in GRAB_OT1.0_ expressing paraventricular and ventral tegmental neuron. Note isolated release sites indicated by pink arrows in **B**. (**C**) Average number of release sites of paraventricular and ventral tegmental neurons (PVN: 8.5±1.9, *n* = 10 neurons; VTA: 6.3±0.8; *n* = 10 neurons; *U* = 42.0, *p* = 0.569). (**D**) Pixel-wise maximal *Δ*F/F_0_ at the isolated release site indicated by the pink arrows in **B**. Fitting the data points with a single-exponential decay function (pink line) yields an estimated oxytocin spatial spread length constant of 0.79 μm at PVN and 0.76 μm at VTA. (**E**) Summary plot of oxytocin diffusion curves from putative single release sites gives an average spatial spread length constant of for oxytocin at paraventricular and ventral tegmental neurons (PVN: 0.78±0.02 μm, *n* = 25 from 10 neurons; VTA: 0.75±0.03, *n* = 21 from 8 neurons; *U* = 281.0, *p* = 0.691). Note the average diffusion curve in black. (**F**) Ten *Δ*F/F_0_ responses evoked by single-pulse stimuli at the single release site (indicated by the pink arrows in **B**) of GRAB_OT1.0_ expressing paraventricular and ventral tegmental neurons. (**G**) Amplitude histograms of *Δ*F/F_0_ responses showed nearly equal peaks for paraventricular neurons and multiple, nearly equally spaced peaks for ventral tegmental neurons. (**H-J**) Values for the average (PVN: 1.30±0.15, *n* = 10; VTA: 1.85±0.19, *n* = 10; *U* = 76, *p* < 0.05) and maximal (PVN: 1.5±0.22, *n* = 10; VTA: 2.60±0.34, *n* = 10; *U* = 79.0, *p* < 0.05) vesicular quanta, release probability (PVN: 100.0±0.0%, *n* = 10; VTA: 100.0±0.0%, *n* = 10; *U* = 50.0, *p* < 0.001), and quantal size (PVN: 14.5±2.2 %*Δ*F/F_0_, *n* = 10; VTA: 11.5±1.8 % *Δ*F/F_0_, *n* = 10; *U* = 41.0, *p* = 0.52) at the single release site for paraventricular and ventral tegmental neurons. **(K)** *Δ*F/F_0_ responses of GRAB_OT1.0_ expressing paraventricular and ventral tegmental neuron evoked by local electrical stimuli of 1,920 pulses at 16 Hz. **(L**) Plots of cumulative *Δ*F/F_0_ responses against the stimulus number in GRAB_OT1.0_ expressing paraventricular and ventral tegmental neurons. Note the readily releasable pool size and refilling rate inferred from the *y*-axis intercept and slope of linear fit to late points of the cumulative trace, respectively. (**M-N**) Values for the vesicular refilling rate (PVN: 1.9±0.5 vesicles/per 100 pulses, *n* = 11; VTA: 0.11±0.08 vesicles/per 100 pulses, *n* = 12; *U* = 4.0, *p* < 0.001) and readily releasable pool (PVN: 26.4±5.8, *n* = 4; VTA: 35.4±5.3, *n* = 12; *U* = 85.0, *p* = 0.255 for whole cells; PVN: 5.0±1.9, *n* = 10; VTA: 6.5±0.8; *n* = 12; *U* = 71.0, *p* = 0.121 for single release sites) of GRABOT1.0 expressing paraventricular and ventral tegmental neurons.

Likewise, we examined a genetically encoded peptide sensor for orexin, OxLight1 (Duffet et al., 2022), in the mouse nucleus accumbens. GESIAP showed orexin release and measured an orexin spread length constant of ~0.70 μm at accumbens neurons (**Fig S11**). Together, these results suggest that transmissions mediated by acetylcholine, monoamines, and neuropeptides all share the characteristics of restricted non-volume synaptic transmission. As controls, using the same *ex vivo* approach, we measured the glutamate spread length constant in the mouse amygdala and paraventricular nuclei with iGluSnFR (Marvin et al., 2013) and iGluSnFR3 (Aggarwal et al., 2022), respectively. GESIAP yielded the same glutamate spread length constant of ~0.60 μm at amygdalar and paraventricular neurons (**Figs S12–13**). This slightly smaller spread length constant for glutamate (**Fig S14**) is consistent with previous reports (Jensen et al., 2019; Zhu et al., 2020), as reflects the fact that the negatively charged glutamate is electrophoretically influenceable by excitatory currents (Sylantyev et al., 2008). Collectively, these results validate the use of GESIAP for nanoscopic visualization of transmission mediated by various transmitters.

## DISCUSSION

This study reports a new image analysis program, GESIAP, that integrates optimized MATLAB-based alignment, deconvolution, baseline correction, denoise and background subtraction algorithms into a single analysis program to provide high-sensitivity, high-resolution visualization of transmission mediated by various transmitters. GESIAP provides quantitative analysis of fundamental synaptic properties, including transmitter spatial diffusion extent, quantal size, quantal content, release probability, pool size, and refilling rate at single release sites. Our experiments validate GESIAP as a versatile program applicable of resolving fundamental synaptic properties of transmission of diverse neurotransmitters at several cell types across multiple animal species. While the analysis consistently supports the notion that neuromodulators use restricted non-volume synaptic transmission as their primary intercellular communication mode, it also unveils a broad spectrum of presynaptic release properties that likely contribute to the diverse actions mediated by diverse neuromodulators across different cells, tissues, organs, and organisms.

### Applicability of GESIAP (and genetically encoded sensors) in visualization of transmission

Our study shows that GESIAP is capable of processing image data collected in genetically encoded sensor-based functional imaging experiments, enabling visualization of transmitter release at the nano- or micro-scale spatiotemporal resolution. The high-resolution visualizations of transmitter release immediately help delineate fundamental synaptic properties of transmission mediated by acetylcholine and monoamines, multiple neuropeptides (e.g., oxytocin and orexin), and glutamate. We demonstrate that GESIAP is generally applicable to resolving transmission at various neuronal and non-neuronal cells in mouse, rat, and human preparations.

These experiments provide the opportunity to evaluate performance of recently engineered genetically encoded sensors, and facilitate determination of signal-to-noise ratio, dynamic range, kinetics, and basal fluorescence level four primary determinants of sensor performance (**Table S1**). The high signal-to-noise ratio of GRAB_5HT1.0_ and GRAB_OT1.0_ facilitated visualization of transmission in every tissue preparation examined. The signal-to-noise ratios of GRAB_NE1m_ and GRAB_DA2.0_ are less than optimal — future sensor versions with higher SNR will improve their performance across preparations. Dynamic range is another key property underlying sensor performance. GRAB_NE1m_, with an affinity closer to endogenous concentrations, performs better than GRAB_NE1h_ Most bacterial periplasmic binding protein-based sensors (e.g., iAChSnFR and iGluSnFR) and many G protein-coupled receptor-based sensors (except GRAB_ACh2.0_ and GRAB_ACh3.0_) have kinetics fast enough to resolve temporal properties of slow-acting neuromodulatory transmission. While most transmitter sensors have basal fluorescence levels sufficient for identification of expressing cells, a few sensors, such as dLight1 and OxLight1, would benefit from higher basal fluorescence.

### Implications of GESIAP in understanding neuromodulation

GESIAP supports a model of spatially restricted synaptic transmission as a key conserved intercellular communication mode for slow-acting transmitters. The widely accepted volume transmission theory of intercellular neuromodulatory communication purports that acetylcholine and monoamines diffuse into local areas, affecting many types of nearby cells; and neuropeptides are thought to travel even farther, influencing local cells as well as distant cells millimeters away (Agnati et al., 1992; Borroto-Escuela et al., 2018). However, direct visualization of transmitter spatial diffusion with GESIAP shows that acetylcholine, monoamines, and multiple neuropeptides all use highly restricted synaptic transmission as a primary mode for intercellular communication. Importantly, high-speed two-photon imaging supports this model for acetylcholine, giving a spread length constant of ~0.75 μm and showing independent releases at neighboring acetylcholine release sites in intact brains ((Kazemipour et al., 2019; Borden et al., 2020); our unpublished data). These results support the view that fast- and slow-acting neurotransmitters share conserved, biophysically and economically advantageous synaptic organizational arrangements to restrict transmitter diffusion (Jensen et al., 2019; Zhu et al., 2020; Lin et al., 2021). We note that some transmitters may spread over larger surface areas of postsynaptic cells by having more release sites (e.g., acetylcholine at astrocytes (**Fig 5**) and norepinephrine at multiple types of neurons (**Fig 6**; our unpublished data). Of course, as with fast-acting neurotransmitters glutamate (e.g., via NMDA receptors) and GABA (e.g., via δ subunit-containing GABA_A_ receptors), a small number of neuromodulatory transmitters may escape synaptic areas and diffuse further away to activate high affinity receptors, serving as a complementary mode under certain physiological and pathological conditions (Barbour and Hausser, 1997; Jensen et al., 2019; Borden et al., 2020; Zhu et al., 2020; Aggarwal et al., 2022).

Interestingly, GESIAP reveals a quite broad spectrum of presynaptic release properties. For example, distinct transmitters exhibit highly varied release probabilities, ranging from <5% (e.g., norepinephrine) to 70-100% (e.g., serotonin and oxytocin) at single release sites. Moreover, distinct transmitters display different vesicular quantal contexts, release pool sizes, and refilling rates. Finally, the same transmitters may employ different release properties when acting on different cell types (e.g., acetylcholine at neurons vs. astrocytes), as well as altered vesicular quantal contexts, release pool sizes, and refilling rates to regulate release at different cell targets (e.g., oxytocin at paraventricular vs. ventral tegmental neurons). These results implicate wide-ranging synaptic properties in diversifying neuromodulatory actions and validate GESIAP as a versatile program for decoding regulations of intercellular transmission.

### Implications of GESIAP in decoding behaviors and diseases

In this study, we validate GESIAP as an effective method to resolve fundamental properties of synaptic transmission. These properties define the precision (e.g., spatial diffusion extent), strength (e.g., quantal size and quantal content), sustainability (e.g., vesicular refilling rate), and short- and long-term plasticity (e.g., release probability) of intercellular communication (Kaeser and Regehr, 2014; Jackman and Regehr, 2017). These processes underpin countless physiological and behavioral actions, and dysregulated properties are indicative of pathological conditions (Greengard, 2001; Sudhof, 2018). Electrophysiologists have made strenuous efforts to perform numerous painstaking experiments to resolve these parameters in various glutamatergic and GABAergic synapses (von Gersdorff and Borst, 2002; Wu et al., 2014; Pulido and Marty, 2017). Here we show that, with GESIAP, a few simple sets of genetically encoded sensor-based functional imaging experiments effectively measure all these parameters.

Delineating the key regulatory parameters of transmission also facilitates inference of how neuromodulatory transmission functions across the brain and body (Greengard, 2001; Sudhof, 2018). Although patch-clamp electrophysiology has delineated much of our understanding of fast-acting glutamatergic and GABAergic transmission, it is ineffective at synaptic transmissions mediated by minimal and/or desensitizing neuromodulator-induced electrophysiological responses (Dani and Bertrand, 2007; Nadim and Bucher, 2014). Alternative methods, like microdialysis and voltammetry, despite many recent improvements, still lack sufficient spatial and/or temporal resolution to decipher properties of transmission (Olive et al., 2000; Robinson et al., 2008; Darvesh et al., 2011; Li et al., 2022). Thus, little is known about the precise regulation and exact function of intercellular communication mediated by neuromodulators. For example, prior work has established the essential role of cholinergic transmission in initiating and maintaining wakefulness and attention critical for behavior (Picciotto et al., 2012; Ballinger et al., 2016). Adrenergic transmission has also been implicated in wakefulness and attention (Carter et al., 2010; Bari et al., 2020; Breton-Provencher et al., 2022), yet its exact functional role remains mysterious since ablating adrenergic neurons does not abolish arousal and attention-dependent behaviors (Aston-Jones and Cohen, 2005; Scammell et al., 2017). Here, we use GESIAP to reveal for the first time that adrenergic transmission exhibits an unusually low release probability, strong facilitation, and prolonged asynchronous release, and it releases norepinephrine via many release sites (**Fig 6**; our unpublished data), distinguishing it from transmission mediated by all other neurotransmitters. These adrenergic synaptic properties predict a unique linear, fine-tuned input-output information computational scheme that is more energy-consuming and vulnerable to system runaway than other transmitters (Kaeser and Regehr, 2014; Jackman and Regehr, 2017). This suggests a special, potentially vital role for adrenergic transmission in regulation of precision- and attention-demanding behavior. This is a testable hypothesis that will be exciting to explore. Similarly, we expect GESIAP to enable resolution of fundamental transmission properties of other monoaminergic transmitters and hundreds of neuropeptidergic transmitters, which should pave the way to understanding their regulations and roles in brain and body actions.

GESIAP will aid the study of disease etiopathology and development of effective therapeutic interventions. For example, based on the finding of diminishing acetylcholine release and deteriorating cholinergic neurons in Alzheimer’s brains (Mash et al., 1985), FDA approved Alzheimer’s drugs that directly or indirectly inhibit acetylcholinesterase to boost cholinergic transmission. Targeting acetylcholine after its release is a poor therapeutic strategy due to lack of appreciation of the intact acetylcholinesterase activity in Alzheimer’s brains and precise spatiotemporal regulation of cholinergic transmission (Borden et al., 2020; Zhu et al., 2020). This may partially explain why such medicines have limited efficacy in cognitive improvement, due to interference with transmission precision (cf. (Barbour and Hausser, 1997; Sarter et al., 2009)), and upon medication termination, they induce irreversible, accelerated deterioration, due to homeostatic regulation (Zemek et al., 2014; Ashford, 2015). GESIAP can define regulatory parameters of cholinergic transmission, thereby setting a crucial baseline for detecting specific defects in Alzheimer’s cholinergic transmission (e.g., those mediated by Tau mutants affecting presynaptic release machinery; cf. (Tracy et al., 2022)) and for restoring healthy transmission by developing therapeutics targeting presynaptic release. Because acetylcholine, monoamines, and neuropeptides are involved in a plethora of physiological actions and numerous neurological and psychiatric disorders (Ballinger et al., 2016; Nusbaum et al., 2017; Okaty et al., 2019; Volkow et al., 2019), we expect GESIAP to become a major player in basic and clinical neuromodulatory research in the near future.

## METHODS

### Animal and human preparations

Male and female C57BL/6J mice and Sprague Dawley rats were used in this study. Animals were maintained in the animal facility at the University of Virginia and family- or pair-housed in the temperature-controlled animal room with 12-h/12-h light/dark cycle. Food and water were available *ad libitum*. Primary human dermal fibroblasts established from skin biopsies from de-identified healthy human donors were transduced to neuron-competent fibroblasts using a previously established protocol (Vadodaria et al., 2016). Serotonergic fibroblasts, capable of expressing serotonergic and neural transcription factors were seeded on tissue culture grade plastic petri dishes/slides/plates and, after 24 hours, the medium was changed to induced neuron conversion medium over 3 weeks. The induced serotonergic neurons were then switched to neural maturation medium to mature over 4-8 weeks. Genetically encoded sensors were expressed in human cultured neurons for ~18 hours using Sindbis virus prior to imaging experiments. All procedures for animal and human cell line experiments were performed following protocols approved by the Animal Care & Use Committee of the University of Virginia and in accordance with US National Institutes of Health guidelines.

### Acute brain slice preparations

Acute geniculate, raphe, hippocampal, entorhinal cortical, amygdalar, striatal, and hypothalamic, as well as pancreas and adrenal tissue slices were prepared from postnatal (P)25-60 animals deeply anesthetized by xylazine-ketamine as described in our previous reports (Wang et al., 2015b; Zhang et al., 2018). Animals were decapitated and the brain block containing the lateral geniculate nucleus, dorsal raphe, hippocampus, entorhinal cortex, amygdala, striatum, or paraventricular nucleus was quickly removed, and the pancreas or adrenal gland was quickly dissected, before placing into cold (0-4°C) oxygenated physiological solution containing (in mM): 125 NaCl, 2.5 KCl, 1.25 NaH_2_PO_4_, 25 NaHCO_3_, 1 MgCl_2_, 25 dextrose, and 2 CaCl_2_, pH 7.4. Brain blocks were directly sectioned into 400-μm-thick brain tissue slices, whereas the pancreas and adrenal gland were first embedded in low-melting temperature agar (2.5% in PBS) and then sectioned into 400-μm-thick tissue slices, using a DSK microslicer (Ted Pella Inc.). The tissue slices were kept at 37.0 ± 0.5 °C in oxygenated physiological solution for ~0.5-1 hour before imaging. During imaging, the slices were submerged in a chamber and stabilized with a fine nylon net attached to a platinum ring. The recording chamber was perfused with oxygenated physiological solution. The half-time for the bath solution exchange was ~6 s, and the temperature of the bath solution was maintained at 34.0 ± 0.5 °C. All antagonists were bath-applied.

### Sindbis and lentivirus preparation and expression

Genetically encoded glutamate, acetylcholine, monoamine, and neuropeptide fluorescent sensors were subcloned into Sindbis and/or lentiviral constructs, and viral particles were produced following our previous studies (Lim et al., 2017; Jing et al., 2018). In brief, fluorescent sensors and their variants were sub-cloned into Sindbis viral vector pSinREP5 with Xba1 and Sph1 restriction digestion, and into lentiviral vector pLenti-with BamH1 and Xho1/Not1, either under a human synapsin promoter for neuronal expression, or under a GFAP promoter for astrocytic expression.

Expression of genetically encoded fluorescent sensors was performed as previously reported (Lim et al., 2017; Jing et al., 2018). P25-60 animals were initially anesthetized by an intraperitoneal injection of ketamine and xylazine (10 and 2 mg/kg, respectively), and then placed in a stereotaxic frame. A glass pipette was used to penetrate the lateral geniculate nucleus, dorsal raphe, hippocampus, entorhinal cortex, amygdala, striatum, or paraventricular nucleus according to stereotaxic coordinates, or the surgically exposed pancreas and adrenal gland, to deliver ~50 nl of Sindbis or lentiviral solution by pressure injection to infect neurons, astrocytes, or nonneuronal cells. Human cultured neurons were infected by including ~50 nl of viral solution in culture media. Experiments were typically performed within 18 ± 4 hours of Sindbis viral infection or 1-2 weeks after lentiviral infection.

### Immunostaining of human serotonergic neurons

Human fibroblast-derived serotonergic neurons were fixed with 4% paraformaldehyde for 30 minutes at room temperature. Fixed neurons were washed three times with phosphate buffered saline with 0.1% Tween 20, and blocked and permeabilized with a solution containing 2% bovine serum albumin (Rockland Immunochemicals, Pottstown, PA), 1% fish skin gelatin (MilliporeSigma), 0.02% saponin (Alfa Aesar, Haverhill, MA), 15% horse serum (Thermo Fisher, Waltham, MA) in PBS with 0.3% Triton X-100 (MilliporeSigma) for 4 hours at room temperature. Immunostaining was done with overnight incubation of primary antibodies, including goat antiserotonin (1:500; Abcam; Cat No: ab66047) and mouse anti-tubulin β3 (Clone Tuj1; 1:500; Biotechne; Cat No: MAB1195) at 4°C, and then 2 hours incubation of secondary antibodies, including anti-goat AlexaFluor 488 (Abcam; Cat No: Ab150133) and anti-mouse AlexaFluor 594 (Abcam; Cat No: Ab150128) at room temperature. Nuclear staining was done with 4’,6-diamidino-2-phenylindole (DAPI) (1:1000; Thermo Fisher) for 5 minutes at room temperature. Immunostained neurons were washed with PBS and mounted using Vectashield (Vector Laboratories, Burlingame, CA) for imaging.

### Fluorescence imaging

Due to the slow nature of neuromodulatory transmission, long-term imaging (from seconds to minutes) became necessary to capture the action of transmitters. To minimize drift and fluctuation vital for high-resolution visualization of transmitter release-induced fluorescence responses (Lin et al., 2021), a stable recording/stimulation and imaging setup was used to carry out all imaging experiments (Wang et al., 2015a). Wide-field epifluorescence imaging was performed using a Hamamatsu ORCA FLASH4.0 camera (Hamamatsu Photonics, Japan), and fluorescent sensor expressing cells in acutely prepared tissue slices were excited by a 460-nm ultrahigh-power low-noise LED (Prizmatix, Givat-Shmuel, Israel) (Wang et al., 2015a; Jing et al., 2018). Fluorescence signals were collected with an Olympus 40x water-immersion objective with a numerical aperture of 0.8. The microscopic point-spread function on our image setup was measured with 23 nm green GATTA beads (GATTAquant GmbH, Hiltpoltstein, Germany) (**Fig S2**). The frame rate of the FLASH4.0 camera was set to 10-50 Hz. To synchronize image capture with electrical stimulation, the camera was set to external trigger mode and triggered by a custom-written IGOR Pro 6 program (WaveMetrics, Lake Oswego, OR)-based software PEPOI (Wang et al., 2015a). Glutamatergic, cholinergic, monoaminergic, or peptidergic fibers in tissue slices were stimulated with a bipolar electrode placed ~50-200 μm from imaged cells. A single or a train of voltage pulses (400 μs, up to 50 V) to evoke transmitter release. The *Δ*F/F_0_ responses of fluorescent transmitter sensor expressing cells had only weak correlation with the basal fluorescence F_0_ (**Fig S15**), suggesting *Δ*F/F_0_ responses to be largely independent of sensor expression levels.

### Imaging analysis with GESIAP

Fluorescence responses were analyzed with GESIAP, which was created using MATLAB 2021a with MATLAB’s Image Processing Toolbox (Mathworks), and comprise five major algorithmic procedures, including alignment, deconvolution, baseline adjustment, denoise and background subtraction. The alignment procedure utilizes translational intensity-based automatic registration to align images (Reddy and Chatterji, 1996). The alignment minimizes the average pixel displacement of moving images from reference images computed with a nonparametric diffeomorphic image registration algorithm (Vercauteren et al., 2009), and stops at a point of diminishing returns or after 200 iterations. After the translational alignment, GESIAP uses our empirically obtained point spread function in our optical system (Zhu et al., 2020) and Landweber deconvolution algorithms to reverse light distortions caused by blurring. We found that Landweber algorithms performed better than naïve/regularized inverse filtering, Tikhonov regularization, Tikhonov-Miller, and Richardson-Lucy algorithms (Sage et al., 2017). Combining our pre- and post-processing procedures, Landweber algorithms achieved the overall performance comparable to commercial packages, such as Huygens (Ponti et al., 2007). The deconvolution process uses iterative gradient-descent to minimize a least-squares cost function (Landweber, 1951) and, at the same time, imposes a non-negativity constraint at each iteration. We found that 25-50 iterations typically achieved a good trade-off between image improvement and noise amplification, with the latter largely overcome by subsequent denoise. Baseline adjustment compensates for the small fluorescence decay associated with photobleaching by applying double-exponential fit on the first and last 10 seconds of the recordings. Multiplication factors are calculated at individual pixels after independent fitting to produce new flat baselines, which are then applied to the entire image sequence. Since genetically encoded transmitter sensors bind transmitters similar to postsynaptic transmitter receptors, we tested three model synaptic functions, including simple rise and single-exponential synaptic function, alpha synaptic function (Rall, 1967), and double-exponential synaptic function (Destexhe et al., 1994), to fit on fluorescence responses at individual pixels over time to denoise. The response onset is determined by back-extrapolation of the linear line crossing 40% and 80% response points in the rising phase. While three synaptic functions yielded similar synaptic parameters, the double-exponential synaptic function is selected in the final program owing to its best fitting. Sometimes, a bilateral filter, a non-linear filter (Tomasi and Manduchi, 1998), is applied to mitigate noise prior to synaptic function fitting if needed. Finally, background subtraction adjusts the background level by uniformly subtracting an average fluorescence value estimated from a non-responsive region adjacent to the cell.

### Synaptic transmission property analysis with GESIAP

To visualize individual transmitter release sites and estimate postsynaptic transmitter spatial diffusion extent, the maximal electrically evoked maximal *Δ*F/F_0_ responses at individual pixels over time were plotted to create 3D spatial profiles for individual transmitter release sites. Individual release sites were isolated and identified using the density-based spatial clustering algorithm DBSCAN (Ester et al., 1996). Iterative selection and analysis isolated and identified individual release sites from overlapping clusters. Pixels with the maximal *Δ*F/F_0_ responses in individual release sites were assumed to be the centers of release. Using strategies from the super-resolution localization microscopy analysis (Thompson et al., 2002; Sauer, 2013; Small and Stahlheber, 2014), fluorescence *Δ*F/F_0_ intensity profiles were averaged over multiple exposures, multiple releases and/or multiple directions of transmitter diffusion gradients, and fit with a single-exponential decay function. Fitting was performed at well-isolated release sites, and their decay constants were extracted as spatial spread length constants.

To estimate the quantal properties of individual transmitter releases, 20-pulse trains at low frequency (0.1 Hz) were used to evoked transmitter release, and failures and releases of *Δ*F/F_0_ events at isolated individual release sites were analyzed using a quantal analysis approach (Fatt and Katz, 1952), which yielded vesicle quantal size, quantal content, and release probability. To estimate the other presynaptic release properties, 1,920-pulse trains at high frequency (16 Hz) were employed to exhaust transmitter release, and cumulative release against stimulus number plots were fit with a published synaptic transmission model (Neher, 2015), which revealed refilling rate and vesicular pool sizes. Individual pulse-evoked responses evoked by 16 Hz pulse trains were mathematically calculated accounting the sensor fluorescence decay constant. Increasing stimulation intensity or frequency affected certain synaptic properties but not others ((Zhu et al., 2020); see also (Neher, 2015)). Thus, we varied stimulation intensity (i.e., 5-30 V) and frequency (i.e., 0.1-16 Hz) to resolve different synaptic properties in the experiments.

### Statistical analysis

Statistical results were reported as mean±s.e.m. Animals or cells were randomly assigned into control or experimental groups and investigators were blinded to experimental treatments. Given the negative correlation between the variation and square root of sample number, *n*, the group sample size was typically set to be ~10-25 to optimize the efficiency and power of statistical tests. Statistical significances of the means (*p*<0.05; two sides) were determined using Wilcoxon non-parametric tests for paired samples and Mann-Whitney Rank Sum nonparametric tests for non-paired samples. The data that support the findings of this study are available from the corresponding authors upon request.

## ACKNOWLEDGMENTS

We thank members of the Julius Zhu lab for suggestions and technical support.

## AUTHOR CONTRIBUTIONS

JJZ conceived the concept, and led the project with input from ACK, LG, LL, BJV, JSM, SMC, MMZ, LLL, and YL; WSZ, YZ, and REZ developed MATLAB-based image analysis program and analyzed data with assistance from LH and DS; WSZ, YZ, PZ, SG, and LH performed molecular biology experiments and collected imaging data with assistance from KG, MEG, and FA; KCV, FHG, ML, TQ, MJ, JF, JW, ML, and PMB provided key reagents; JJZ and WSZ wrote the manuscript with input from all other coauthors.

---

**Figure S1.**
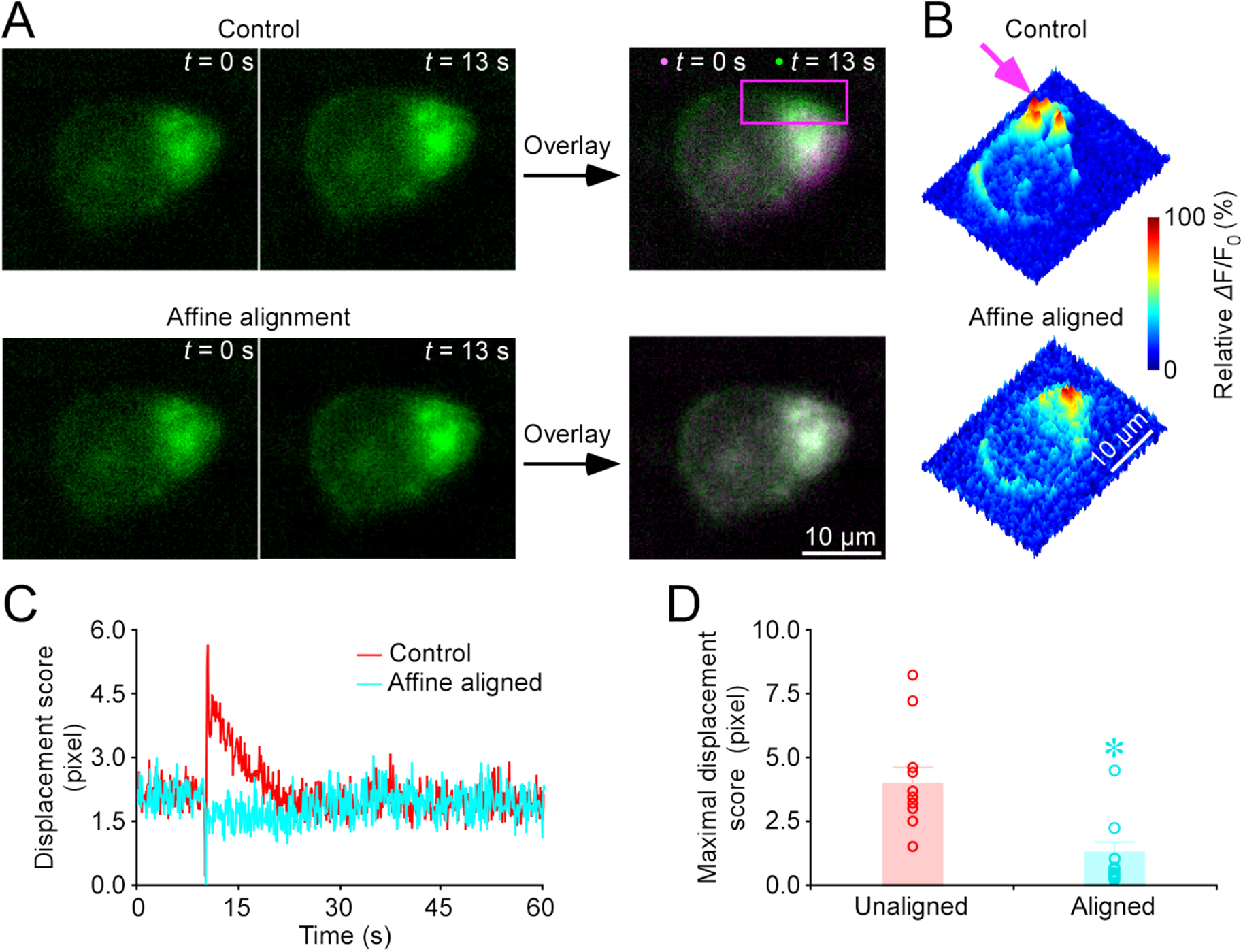
Affine alignment preserves image faithfulness. (**A**) Snapshots and overlay of images captured at two time points of a GRAB_5HT1.0_ expressing geniculate neuron. Note the prominent mismatch in the overlay of control unaligned images. (**B**) 3D spatiotemporal profiling of electrically evoked *Δ*F/F_0_ responses in the GRAB_5HT1.0_ expressing geniculate neuron. Note the spurious “peaks” due to the mismatch (pink arrow indicates one such peak). (**C**) Displacement score of the geniculate neuron. (**D**) Maximal displacement scores before and after translation alignment (*n* = 10 neurons).

**Figure S2.**
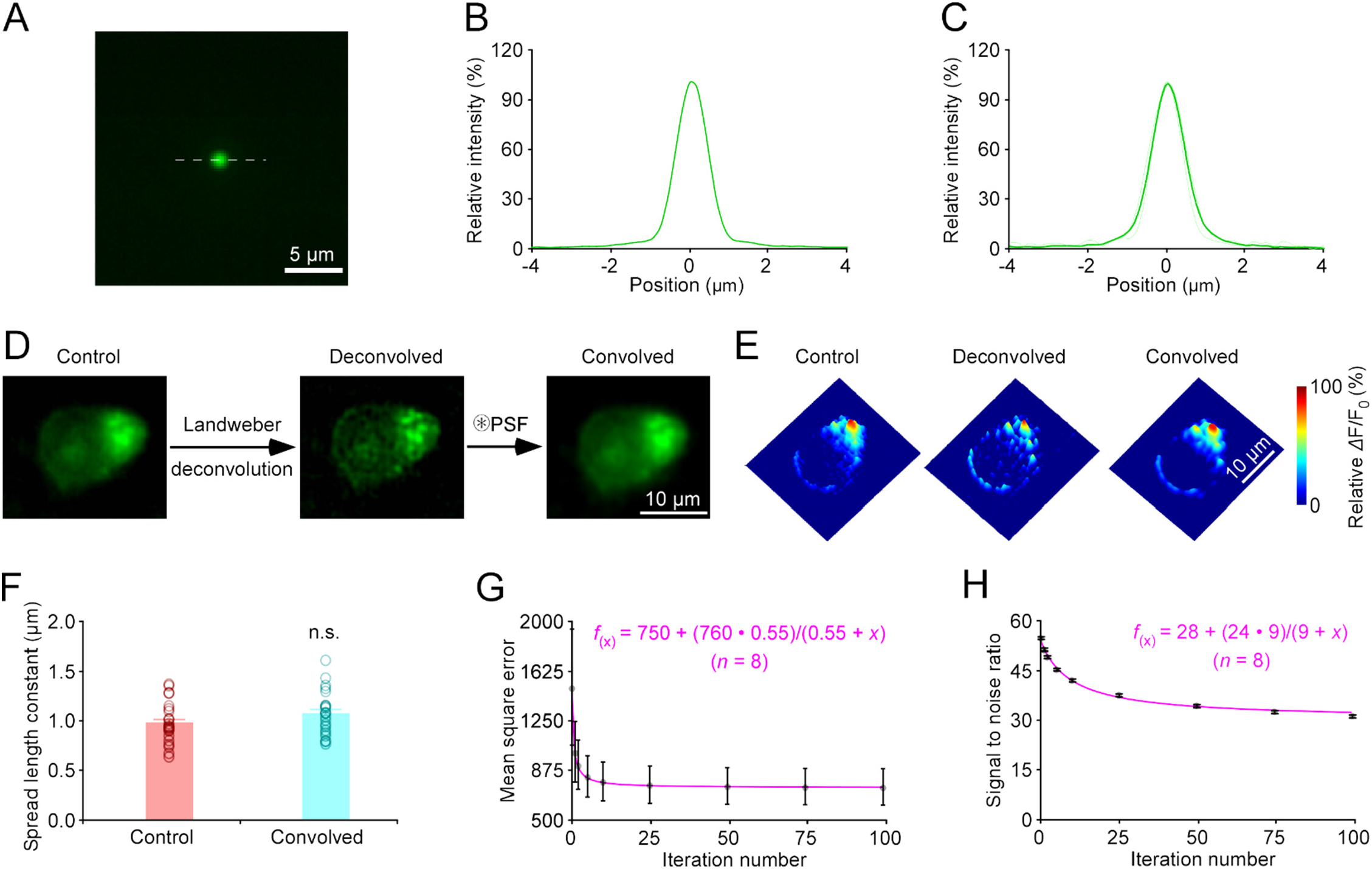
Determine the point-spread function (PSF) for deconvolution. (**A**) Fluorescence image of a 23-nm green GATTA bead under a 0.8 NA 40x objective. (**B**) Point-spread function (PSF) of the 23-nm green GATTA bead shown in (**A**). (**C**) Individual (light green) and average (dark green) PSFs of 23-nm green GATTA beads (*n* = 10). Full width at half maximums (FWHMs) of PSFs of 23-nm green GATTA beads (0.996±0.021 μm, *n* = 10). (**D-E**) Snapshots (**D**), and 3D spatiotemporal profiling (**E**) of electrically evoked fluorescence responses in a GRAB_5HT1.0_ expressing geniculate neuron under control, deconvolved, and convolved processing. (**F**) Average spatial spread length constants for serotonin at geniculate neurons obtained from control and convolved images (control: 1.01±0.06μm, denoised: 1.08±0.05 μm, *Z* = 1.929, *p* = 0.056; *n* = 9 neurons). (**G**) The evolution of mean squared error of control and convolved images during the iterative process. Note the data points fitted with a hyperbolic decay function (pink line; *r^2^* = 0.9999). (**H**) The evolution of signal-to-noise ratio during the iterative process. Note the data points fitted with a hyperbolic decay function (pink line; *r*^2^ = 0.9944).

**Figure S3.**
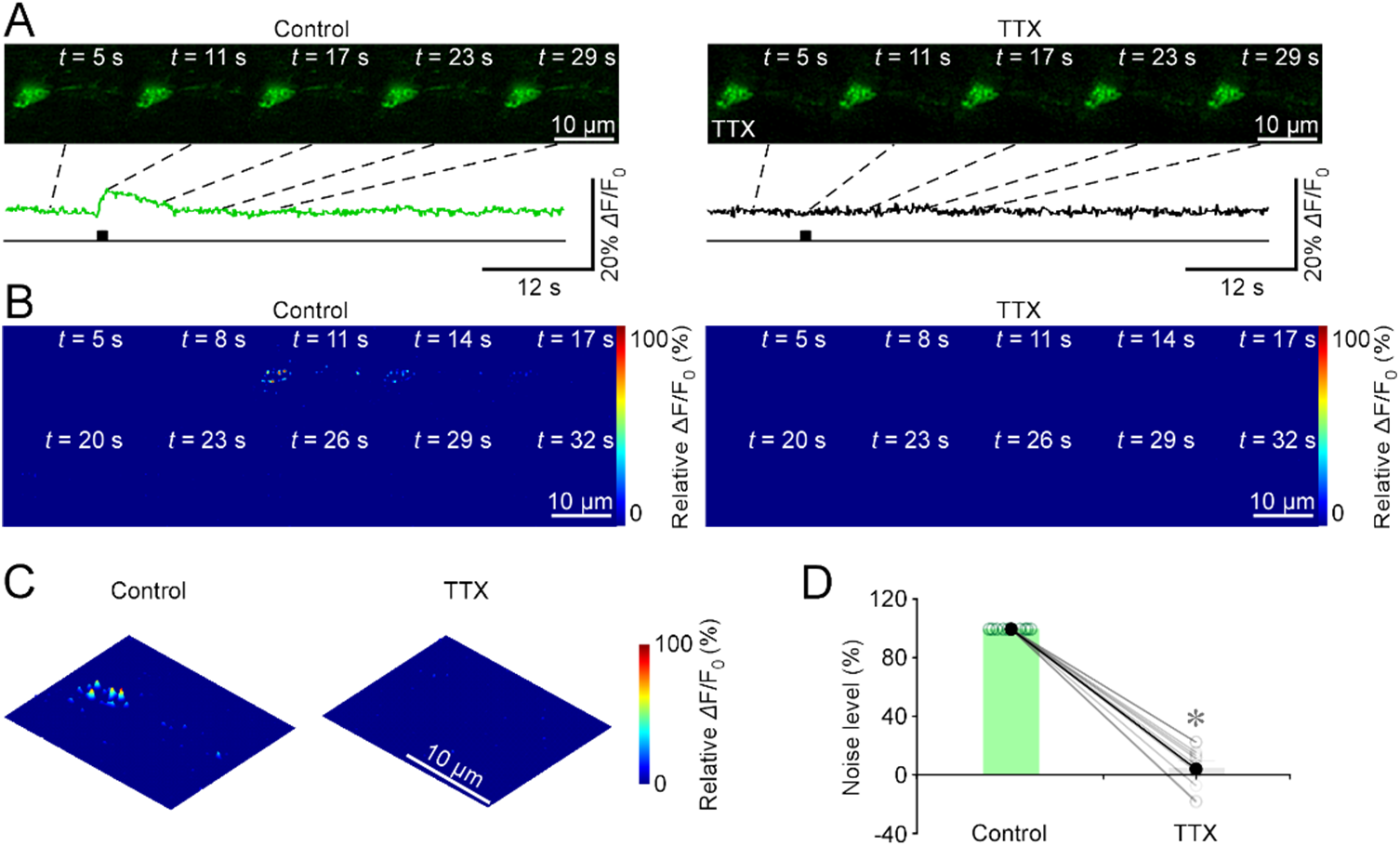
TTX diminishes the evoked *Δ*F/F_0_ responses at GRAB_5HT1.0_ expressing geniculate neurons. (**A-C**) Snapshots (**A**), heatmaps (**B**), and 3D spatiotemporal profiling (**C**) of electrically evoked fluorescence responses in a GRAB_5HT1.0_ expressing geniculate neuron in the normal bath solution and bath solution containing 1 μM TTX. (**D**) Maximal *Δ*F/F_0_ responses under control and TTX conditions (Control: 100.0±0.0% vs. TTX: 5.7±4.9%; *n* = 10 neurons, *Z* = −2.803, *p* = 0.002). Asterisk indicates *p* < 0.01 (Wilcoxon test).

**Figure S4.**
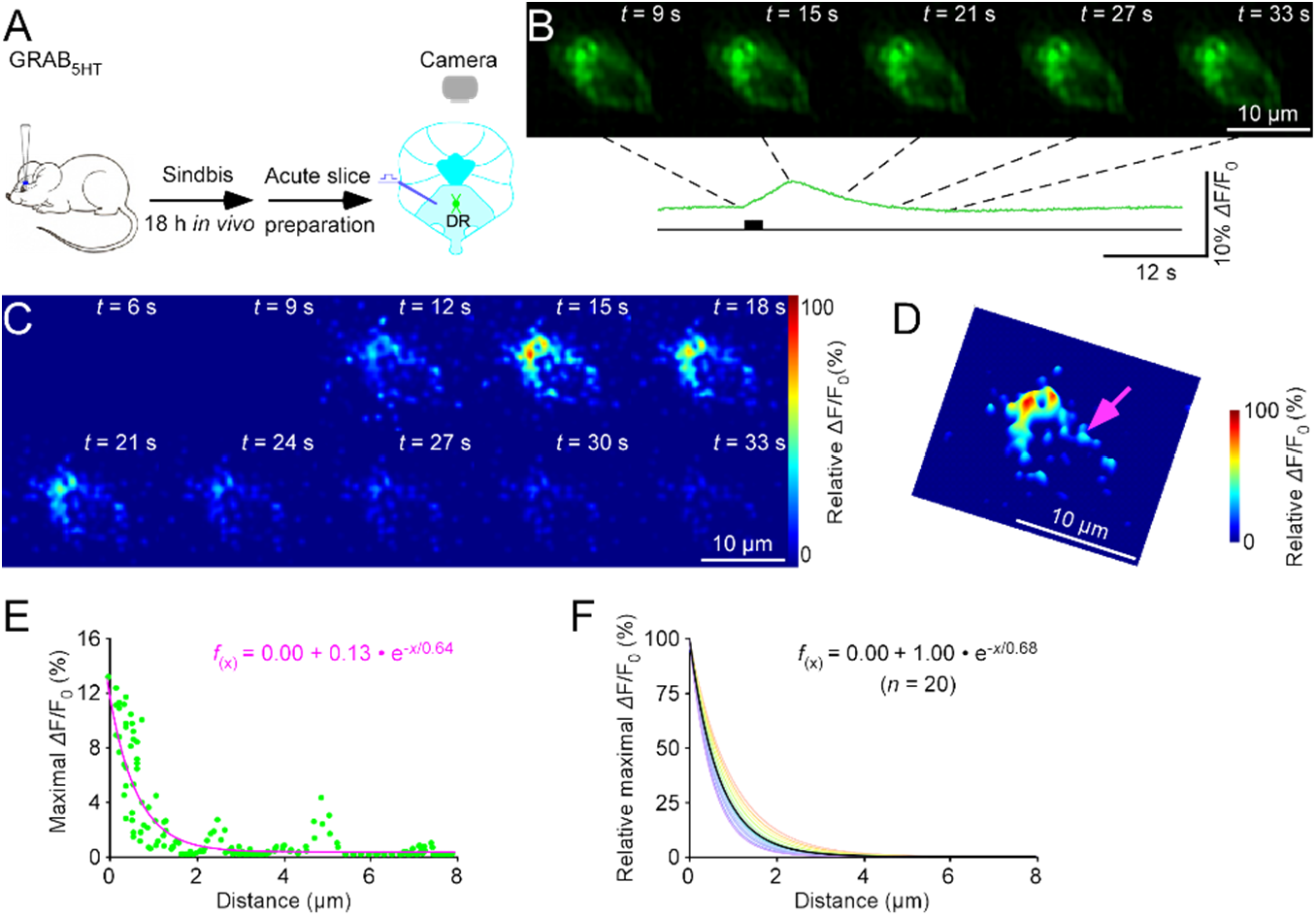
Nanoscopic visualization of serotonergic transmission at raphe neurons. (**A**) Schematic of stimulation-imaging experiment in an *ex vivo* raphe preparation. DR: the dorsal raphe. (**B-D**) Snapshots (**B**), heatmaps (**C**), and 3D spatiotemporal profiling (**D**) of electrically evoked *Δ*F/F_0_ responses in a GRAB_5HT1.0_ expressing raphe neuron. Note the isolated release site indicated by a pink arrow in **D**. (**E**) Pixel-wise maximal *Δ*F/F_0_ at the isolated release site indicated by the pink arrow in **D**. Fitting the data points with a single-exponential decay function (pink line) yields an estimated 5HT spatial spread length constant of 0.61 μm. (**F**) Summary plot of 5HT diffusion curves from putative single release sites gives an average spatial spread length constant of 0.68±0.03 μm for 5HT at raphe neurons (*n* = 20 from 11 neurons). Note the average diffusion curve in black.

**Figure S5.**
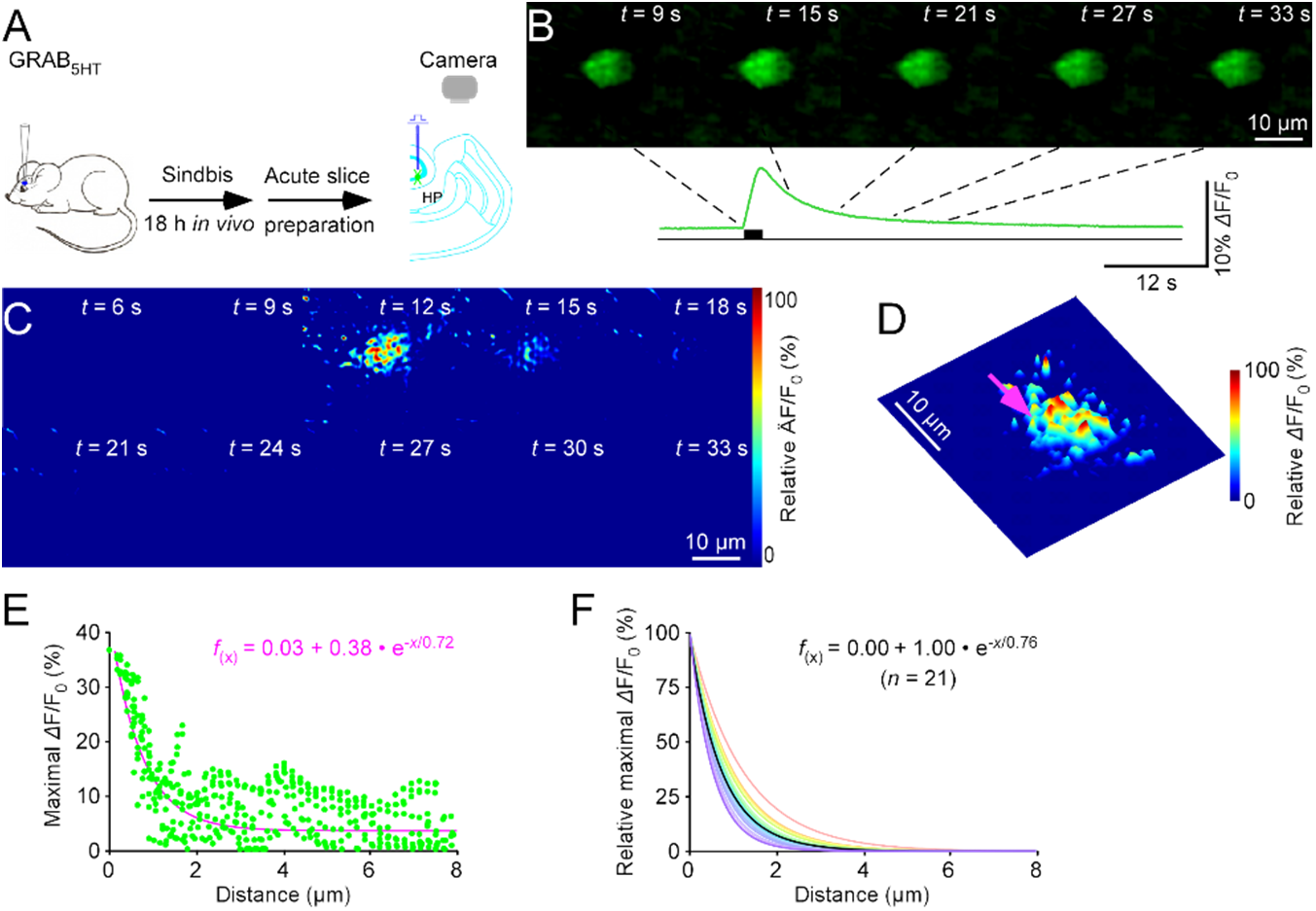
Nanoscopic visualization of serotonergic transmission at rat hippocampal neurons. (**A**) Schematic of stimulation-imaging experiment in an *ex vivo* rat hippocampal preparation. HP: the hippocampus. (**B-D**) Snapshots (**B**), heatmaps (**C**), and 3D spatiotemporal profiling (**D**) of electrically evoked fluorescence responses in a GRAB_5HT1.0_ expressing dentate gyrus neuron. Note the isolated release site indicated by a pink arrow in **D**. (**E**) Pixel-wise maximal *Δ*F/F_0_ at the isolated release site indicated by the pink arrow in **D**. Fitting the data points with a single-exponential decay function (pink line) yields an estimated serotonin spatial spread length constant of 0.67 μm. (**F**) Summary plot of serotonin diffusion curves from putative single release sites gives an average spatial spread length constant of 0.76±0.04 μm for serotonin at dentate gyrus neurons (*n* = 21 from 7 neurons). Note the average diffusion curve in black.

**Figure S6.**
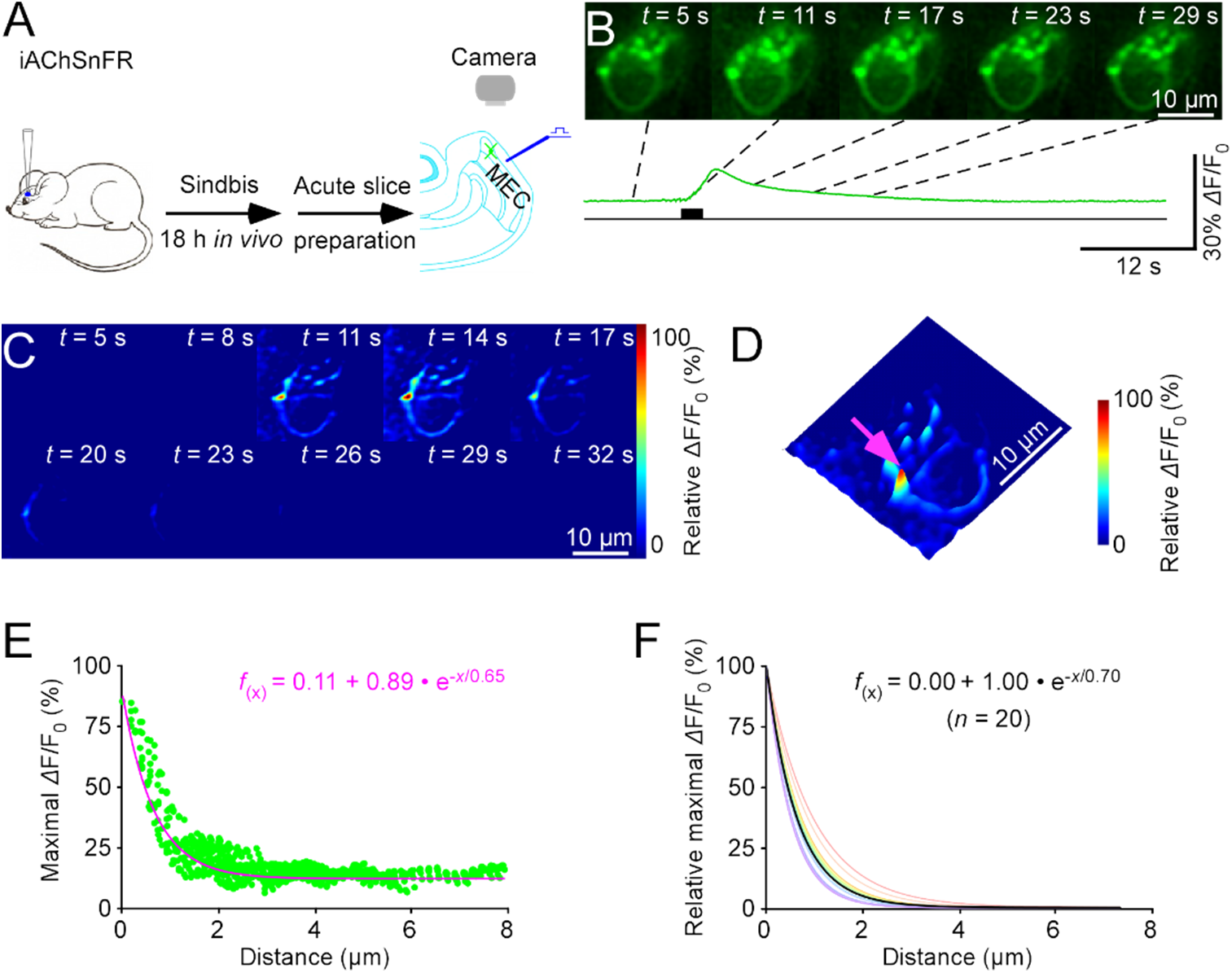
Nanoscopic visualization of cholinergic transmission at entorhinal neurons. (**A**) Schematic of stimulation-imaging experiment in an *ex vivo* entorhinal preparation. MEC: the medial entorhinal cortex. (**B-D**) Snapshots (**B**), heatmaps (**C**), and 3D spatiotemporal profiling (**D**) of electrically evoked fluorescence *Δ*F/F_0_ responses in an iAChSnFR expressing entorhinal stellate neuron. Note the isolated release site indicated by a pink arrow in **D**. (**E**) Pixel-wise maximal *Δ*F/F_0_ at the isolated release site indicated by the pink arrow in **D**. Fitting the data points with a single-exponential decay function (pink line) yields an estimated acetylcholine spatial spread length constant of 0.65 μm. (**F**) Summary plot of acetylcholine diffusion curves from putative single release sites gives an average spatial spread length constant of 0.70±0.03 μm for acetylcholine at entorhinal stellate neurons (*n* = 20 from 10 cells). Note the average diffusion curve in black.

**Figure S7.**
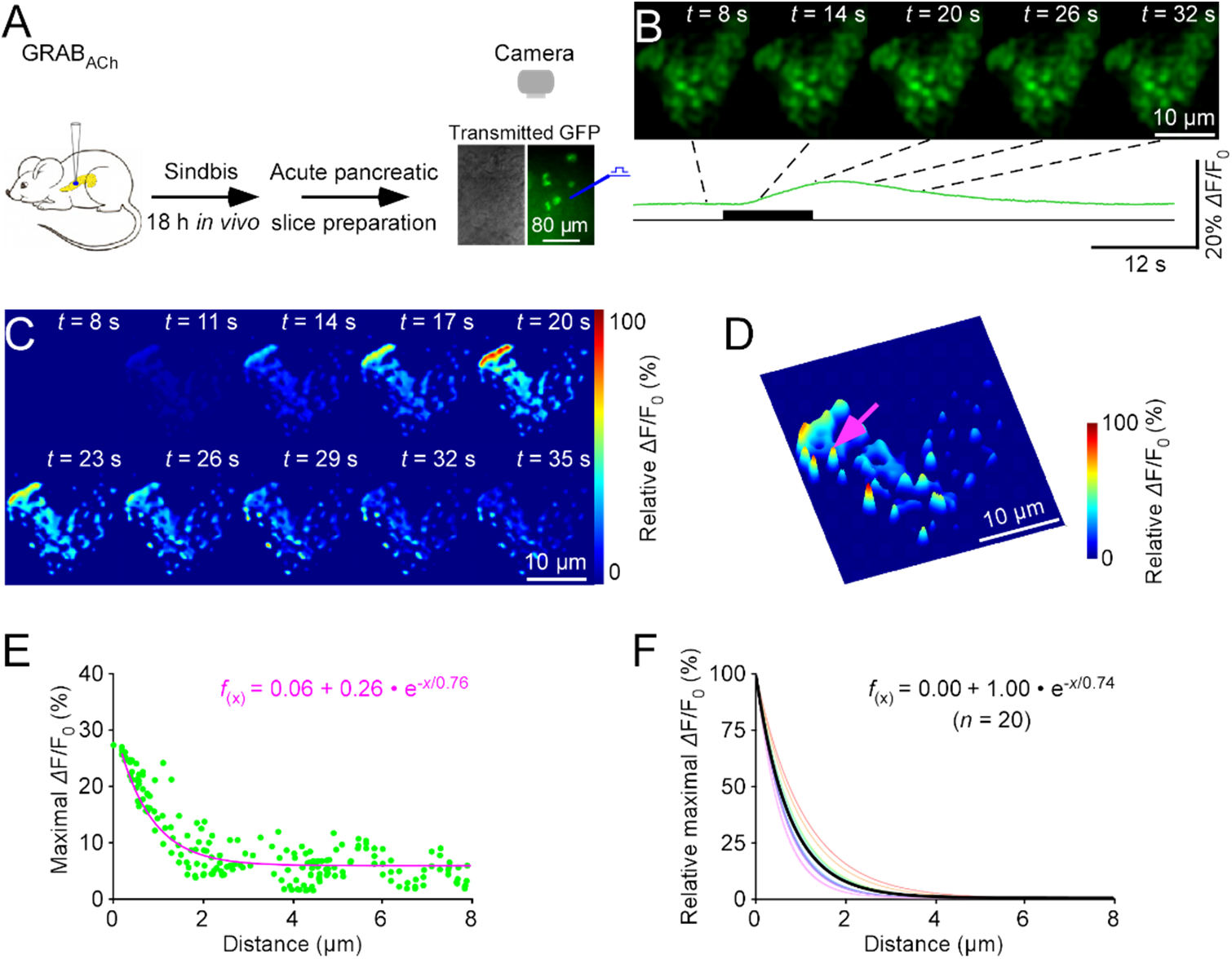
Nanoscopic visualization of cholinergic transmission at pancreatic cells. (**A**) Schematic of stimulation-imaging experiment in an *ex vivo* pancreatic preparation. (**B-D**) Snapshots (**B**), heatmaps (**C**), and 3D spatiotemporal profiling (**D**) of electrically evoked fluorescence responses in a GRAB_ACh_ expressing pancreatic cell. Note the isolated release site indicated by a pink arrow in **D**. (**E**) Pixel-wise maximal *Δ*F/F_0_ at the isolated release site indicated by the pink arrow in **E**. Fitting the data points with a single-exponential decay function (pink line) yields an estimated acetylcholine spatial spread length constant of 0.76 μm. (**F**) Summary plot of acetylcholine diffusion curves from putative single release sites gives an average spatial spread length constant of 0.74±0.03 μm for acetylcholine at pancreatic cells (*n* = 20 from 11 cells). Note the average diffusion curve in black.

**Figure S8.**
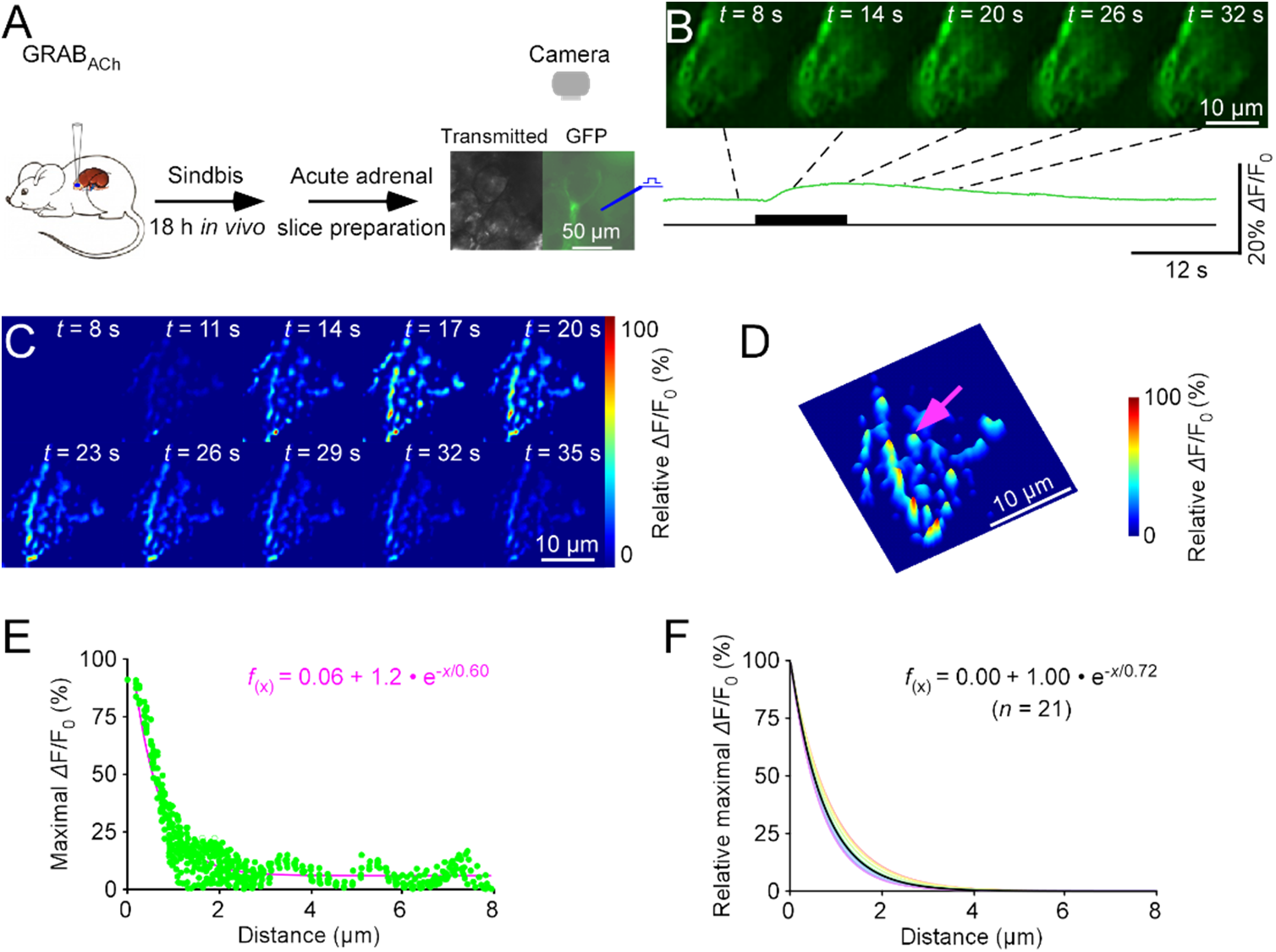
Nanoscopic visualization of cholinergic transmission at adrenal cells. (**A**) Schematic of stimulation-imaging experiment in an *ex vivo* adrenal preparation. (**B-D**) Snapshots (**B**), heatmaps (**C**), and 3D spatiotemporal profiling (**D**) of electrically evoked fluorescence *Δ*F/F_0_ responses in a GRAB_ACh_ expressing adrenal cell. Note the isolated release site indicated by a pink arrow in **D**. (**E**) Pixel-wise maximal *Δ*F/F_0_ at the isolated release site indicated by the pink arrow in **D**. Fitting the data points with a single-exponential decay function (pink line) yields an estimated acetylcholine spatial spread length constant of 0.60 μm. (**F**) Summary plot of acetylcholine diffusion curves from putative single release sites gives an average spatial spread length constant of 0.72±0.03 μm for acetylcholine at adrenal cells (*n* = 21 from 11 cells). Note the average diffusion curve in black.

**Figure S9.**
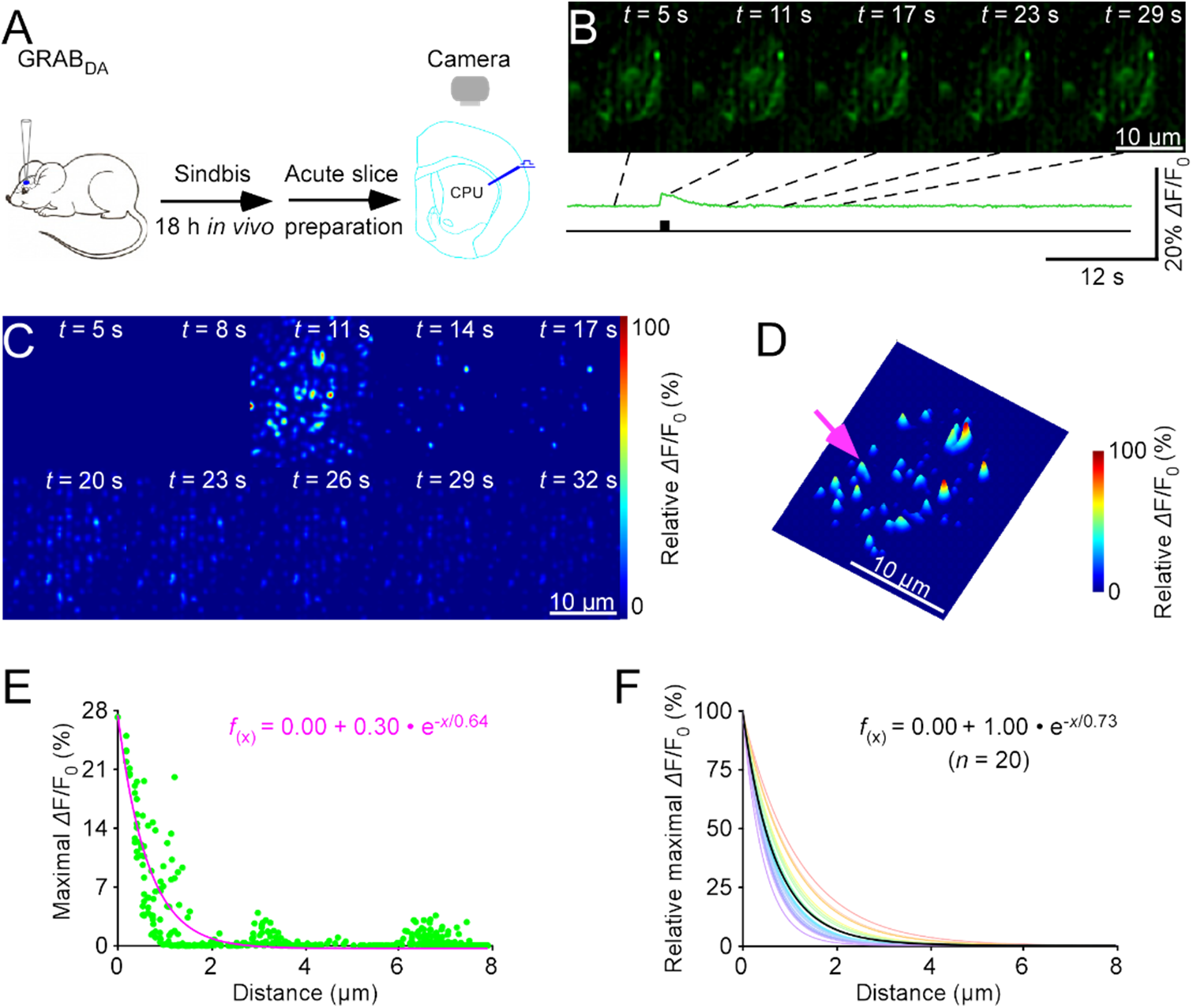
Nanoscopic visualization of dopaminergic transmission at striatal neurons. (**A**) Schematic of stimulation-imaging experiment in an *ex vivo* striatal preparation. DS: the dorsal striatum. (**B-D**) Snapshots (**B**), heatmaps (**C**), and 3D spatiotemporal profiling (**D**) of electrically evoked fluorescence responses in a GRAB_DA2.0_ expressing striatal neuron. Note the isolated release site indicated by a pink arrow in **D**. (**E**) Pixel-wise maximal *Δ*F/F_0_ at the isolated release site indicated by the pink arrow in **D**. Fitting the data points with a single-exponential decay function (pink line) yields an estimated dopamine spatial spread length constant of 0.64 μm. (**F**) Summary plot of dopamine diffusion curves from putative single release sites gives an average spatial spread length constant of 0.70±0.05 μm for dopamine at striatal neurons (*n* = 20 from 10 neurons). Note the average diffusion curve in black.

**Figure S10.**
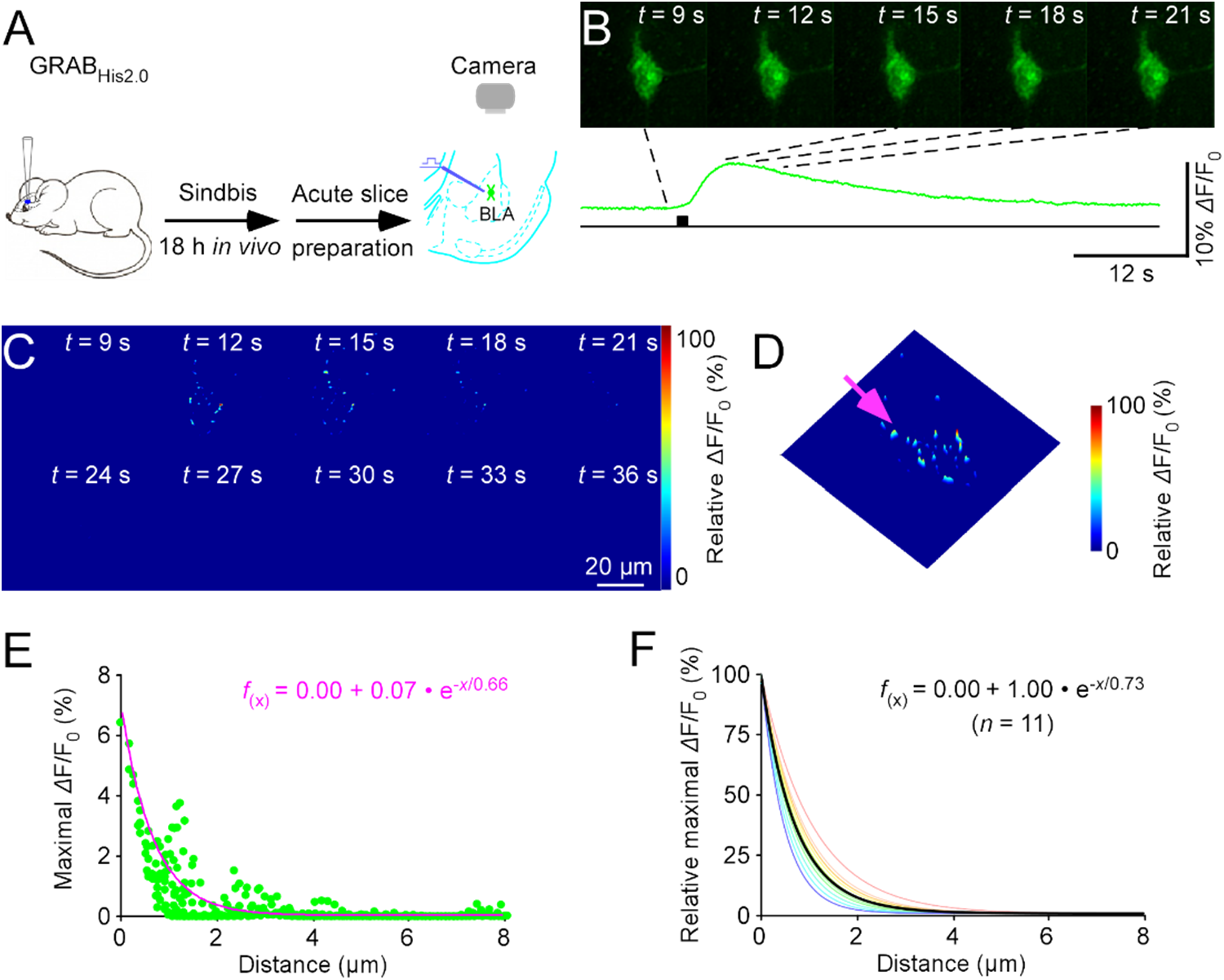
Nanoscopic visualization of histaminergic transmission at amygdalar neurons. (**A**) Schematic of stimulation-imaging experiment in an *ex vivo* amygdalar preparation. BLA: the basolateral amygdala. (**B-D**) Snapshots (**B**), heatmaps (**C**), and 3D spatiotemporal profiling (**D**) of electrically evoked *Δ*F/F_0_ responses in a GRAB_His2.0_ expressing amygdalar neuron. Note the isolated release site indicated by a pink arrow in **D**. (**E**) Pixel-wise maximal *Δ*F/F_0_ at the isolated release site indicated by the pink arrow in **D**. Fitting the data points with a single-exponential decay function (pink line) yields an estimated norepinephrine spatial spread length constant of 0.66 μm. (**G**) Summary plot of norepinephrine diffusion curves from putative single release sites gives an average spatial spread length constant of 0.73±0.05 μm for histamine at amygdalar neurons (*n* = 11 from 6 neurons). Note the average diffusion curve in black.

**Figure S11.**
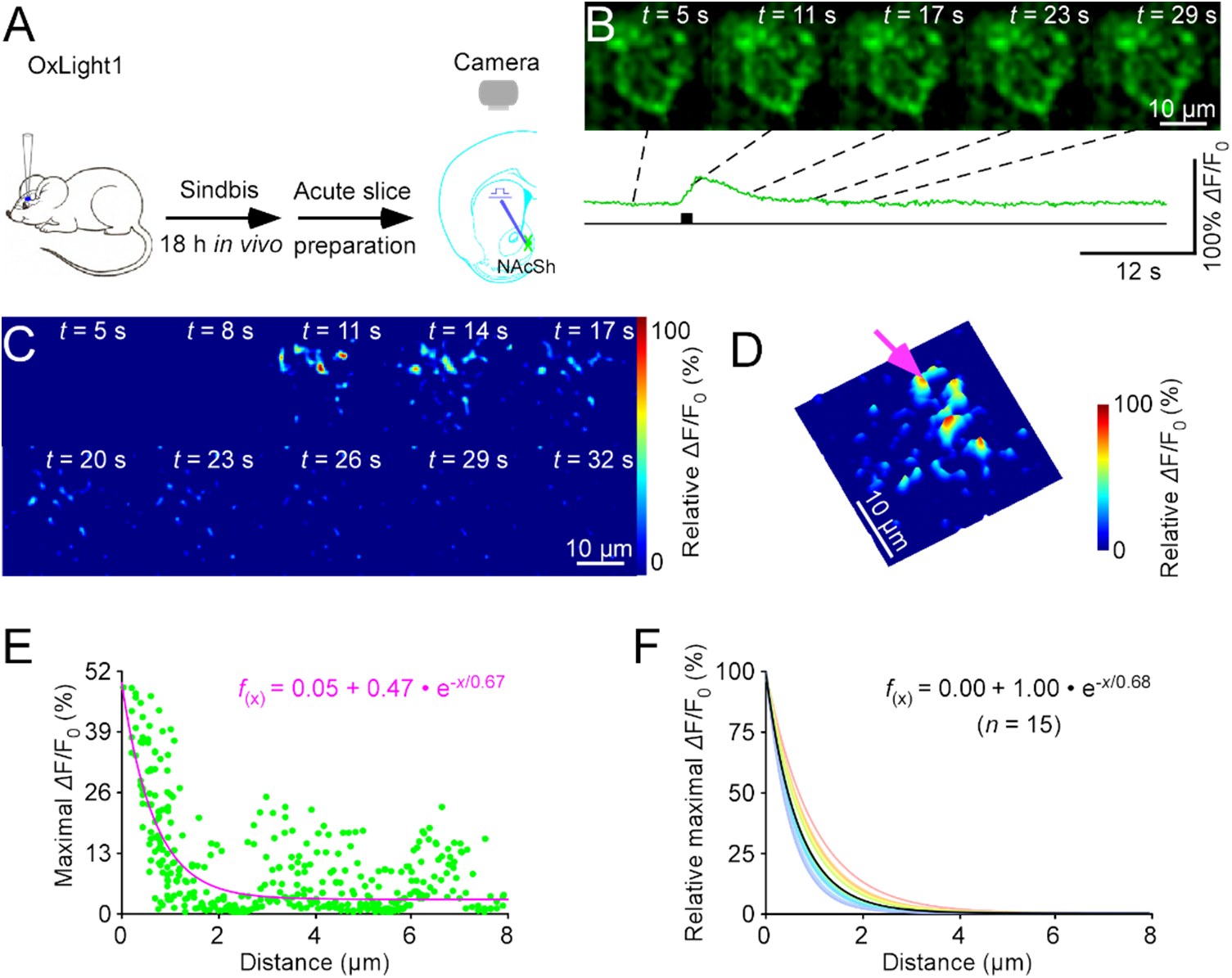
Nanoscopic visualization of orexinergic transmission at accumbens neurons. (**A**) Schematic of stimulation-imaging experiment in an *ex vivo* accumbens preparation. NAcSh: the nucleus accumbens shell. (**B-D**) Snapshots (**B**), heatmaps (**C**), and 3D spatiotemporal profiling (**D**) of electrically evoked *Δ*F/F_0_ responses in an OxLight1 expressing nucleus accumbens neuron. Note the isolated release site indicated by a pink arrow in **D**. (**E**) Pixel-wise maximal *Δ*F/F_0_ at the isolated release site indicated by the pink arrow in **D**. Fitting the data points with a single-exponential decay function (pink line) yields an estimated orexin spatial spread length constant of 0.67 μm. (**G**) Summary plot of orexin diffusion curves from putative single release sites gives an average spatial spread length constant of 0.68±0.04 μm for orexin at nucleus accumbens neurons (*n* = 15 from 6 neurons). Note the average diffusion curve in black.

**Figure S12.**
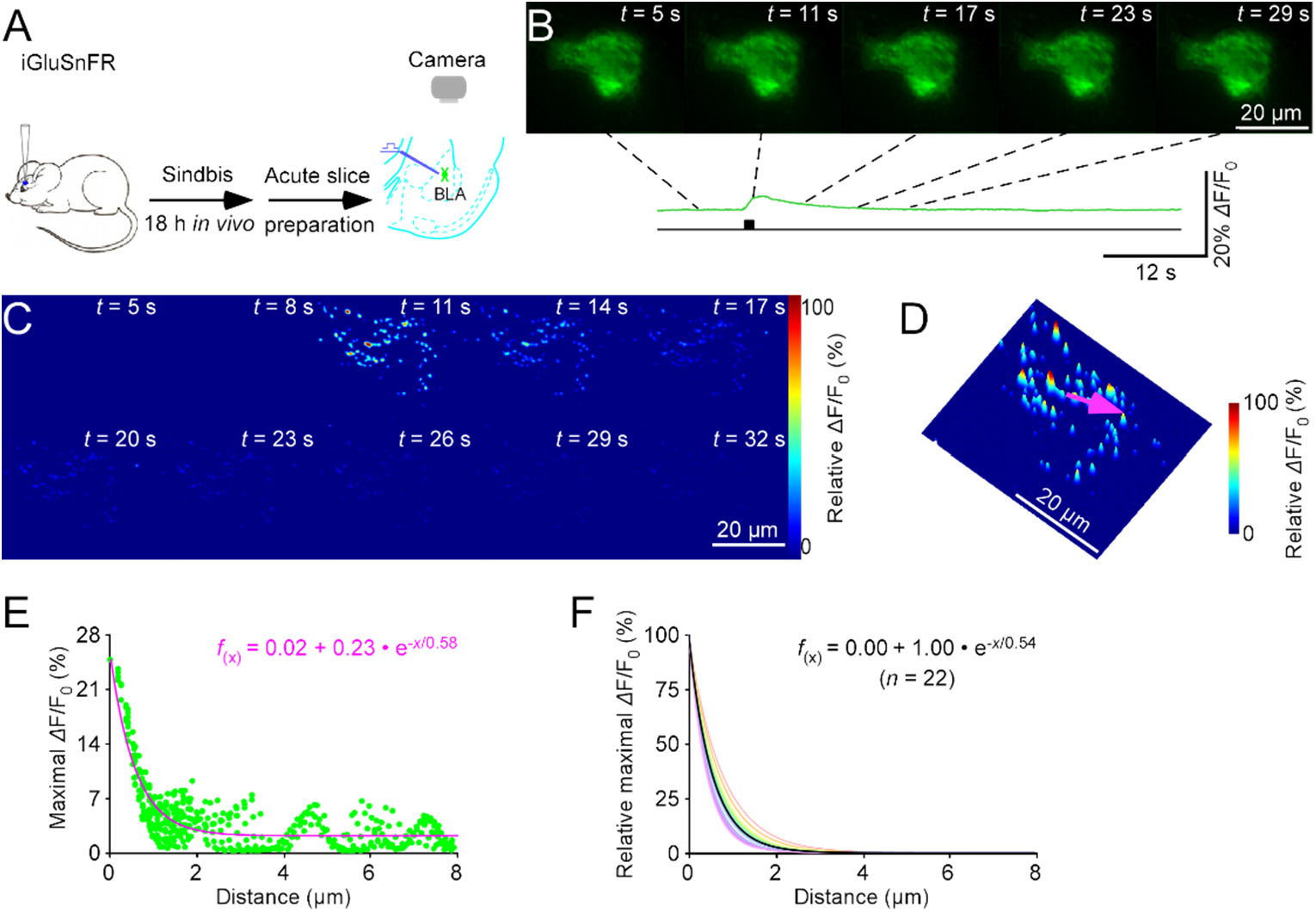
Nanoscopic visualization of glutamatergic transmission at amygdalar neurons. (**A**) Schematic of stimulation-imaging experiment in an *ex vivo* amygdalar preparation. BLA: the basolateral amygdala. (**B-D**) Snapshots (**B**), heatmaps (**C**), and 3D spatiotemporal profiling (**D**) of electrically evoked fluorescence responses in an iGluSnFR expressing amygdalar neuron. Note the isolated release site indicated by a pink arrow in **D**. (**E**) Pixel-wise maximal *Δ*F/F_0_ at the isolated release site indicated by the pink arrow in **D**. Fitting the data points with a single-exponential decay function (pink line) yields an estimated glutamate spatial spread length constant of 0.58 μm. (**F**) Summary plot of glutamate diffusion curves from putative single release sites gives an average spatial spread length constant of 0.54±0.02 μm for glutamate at amygdalar neurons (*n* = 22 from 10 cells). Note the average diffusion curve in black.

**Figure S13.**
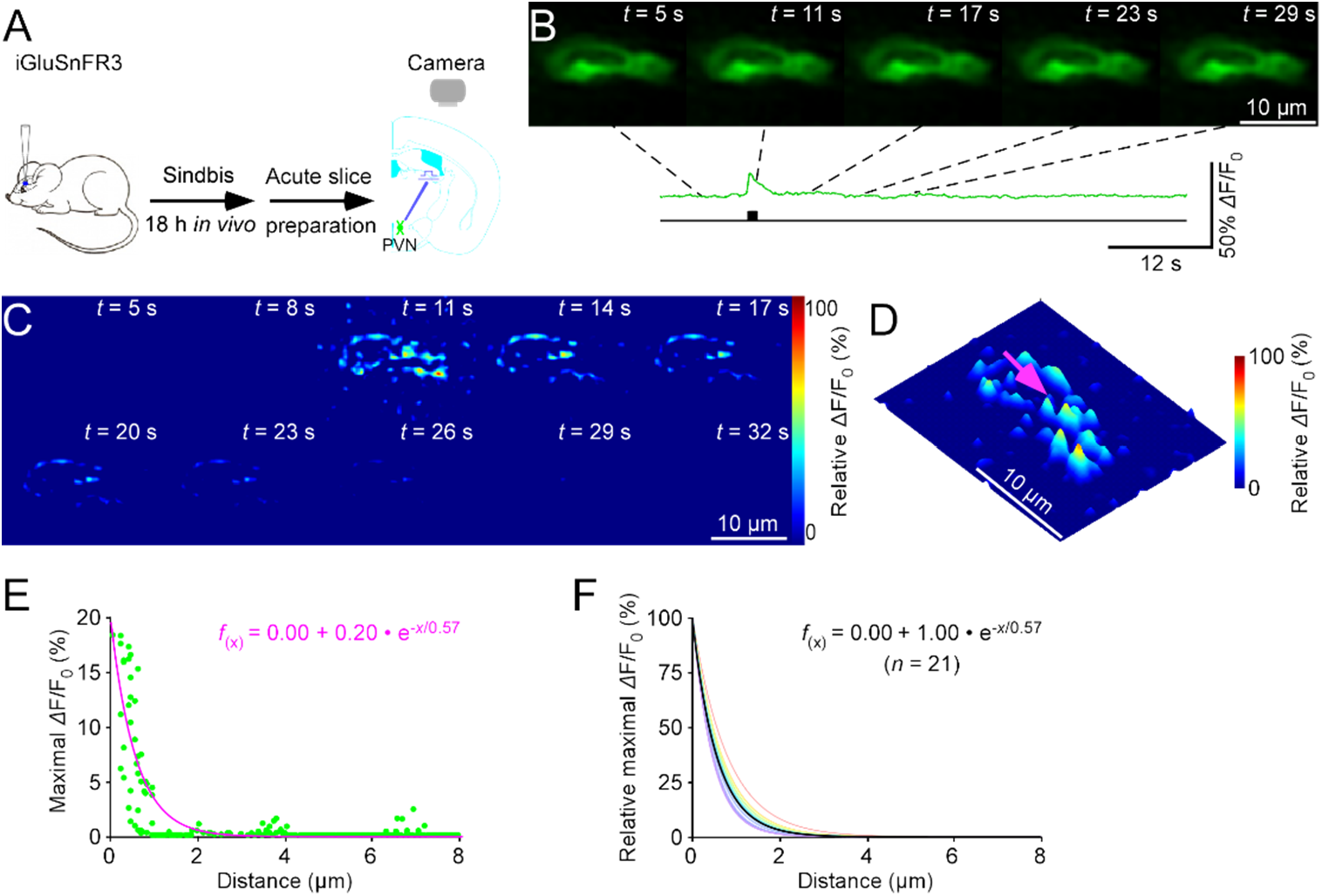
Nanoscopic visualization of glutamate release at paraventricular neurons. (**A**) Schematic of stimulation-imaging experiment in an *ex vivo* paraventricular preparation. PVN: the paraventricular nucleus. (**B-D**) Snapshots (**B**), heatmaps (**C**), and 3D spatiotemporal profiling (**D**) of electrically evoked fluorescence responses in an iGluSnFR3 expressing paraventricular neuron. Note the isolated release site indicated by a pink arrow in **D**. (**E**) Pixel-wise maximal *Δ*F/F_0_ at the isolated release site indicated by the pink arrow in **D**. Fitting the data points with a single-exponential decay function (pink line) yields an estimated glutamate spatial spread length constant of 0.57 μm. (**F**) Summary plot of glutamate diffusion curves from putative single release sites gives an average spatial spread length constant of 0.57±0.02 μm for glutamate at paraventricular neurons (*n* = 21 from 9 cells). Note the average diffusion curve in black.

**Figure S14.**
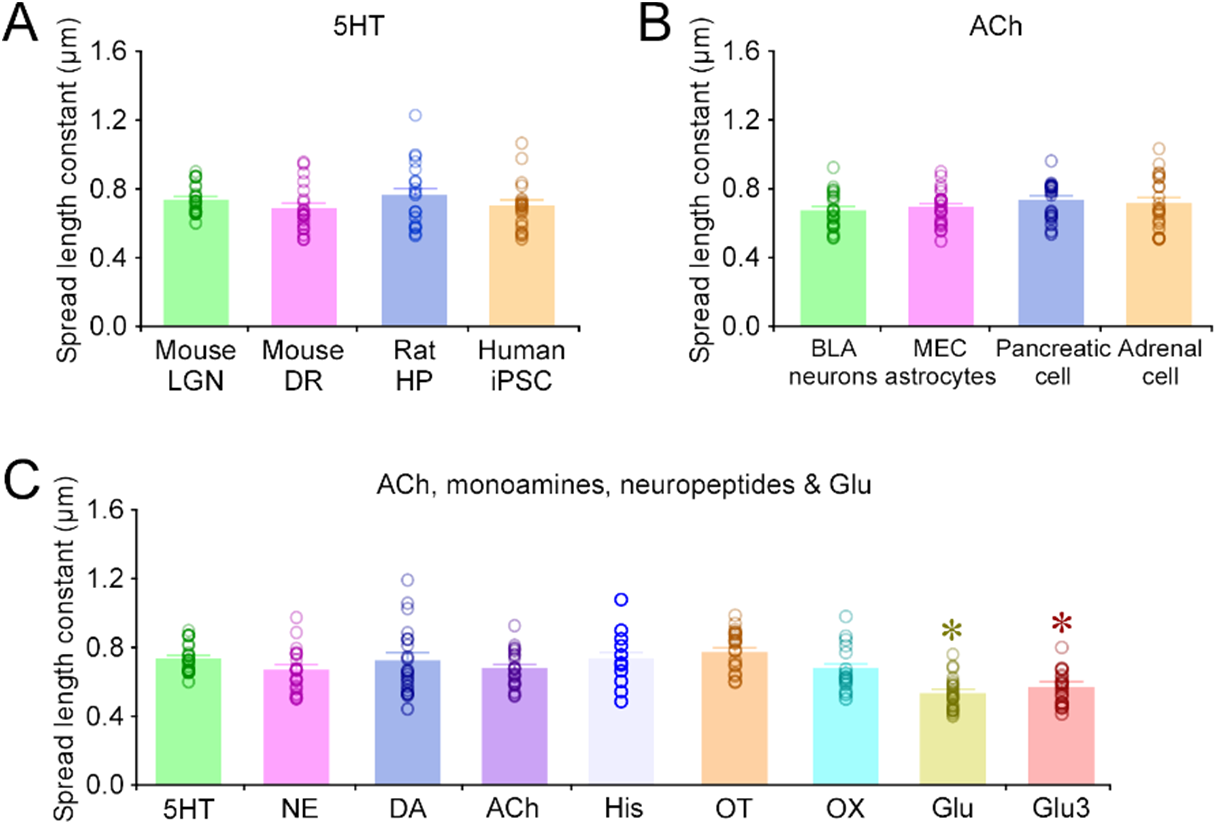
Spread length constants for various transmitters at various cell types of various animals. (**A**) Spread length constants for serotonin (**5HT**) at different nuclei of various animal species, including mouse geniculate neurons (0.74±0.02, *n* = 22), mouse raphe neurons (0.68±0.03, *n* = 20), rat hippocampal neurons (0.76±0.04, *n* = 21), and human fibroblast-derived neurons (0.70±0.03, *n* = 20). Note no differences in the spread length constants for 5HT at different nuclei of various animal species (*p* > 0.05; Mann-Whitney Rank Sum tests). (**B**) Spread length constants for ACh at various mouse cell types, including amygdalar neurons (0.68±0.02, *n* = 21), medial entorhinal astrocytes (0.70±0.02, *n* = 22), pancreatic cells (0.74±0.03, *n* = 20), and adrenal cells (0.72±0.34, *n* = 21). Note no differences in the spread length constants for ACh at various mouse cells (*p* > 0.05; Mann-Whitney Rank Sum tests). (**C**) Spread length constants for various transmitters, including 5HT at mouse geniculate neurons (0.74±0.02, *n* = 22), norepinephrine (**NE**) at amygdala (0.67±0.03, *n* = 20), dopamine (**DA**) at dorsal striatum (0.73±0.05, *n*=20), acetylcholine (**ACh**) at amygdala (0.68±0.02, *n* = 21), histamine (**His**) at amygdala (0.73±0.05, *n* = 11), oxytocin (**OT**) at paraventricular nucleus (0.78±0.03, *n* = 25), orexin (**OX**) at nucleus accumbens (0.68±0.04, *n* = 15), glutamate (**Glu**) at geniculate neurons (0.54±0.02, *n* = 22), and glutamate (**Glu3**) at paraventricular nucleus (0.57±0.03, *n* = 15). Note the significantly smaller spread length constants for glutamate compared to the other transmitters (*p* < 0.05; Mann-Whitney Rank Sum tests).

**Figure S15.**
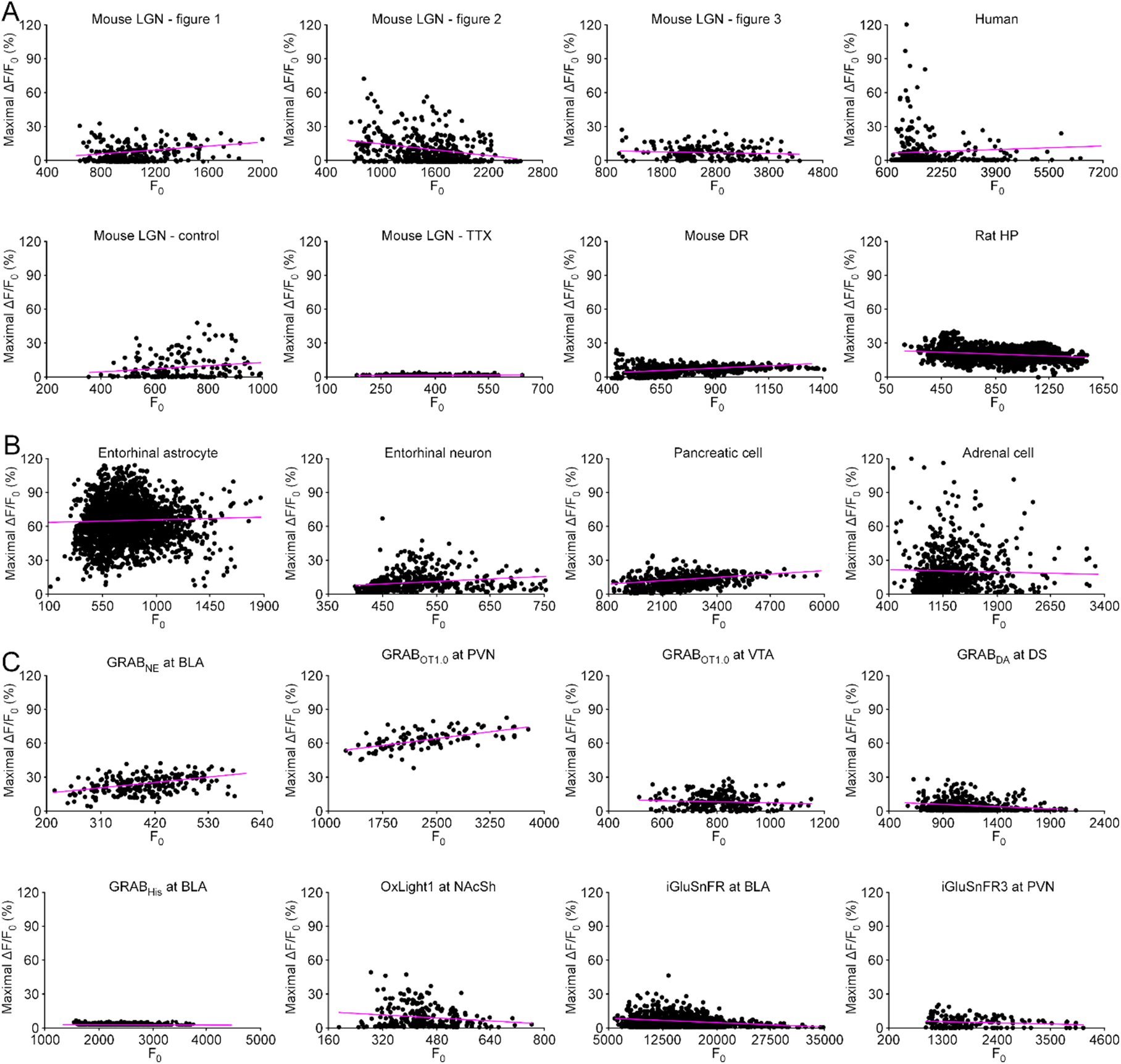
Fluorescence responses are largely independent of sensor expression levels. (**A**) Fluorescence responses are largely independent of GRAB_5HT_ expression. Plots of ΔF/F_0_ against F_0_ of the GRAB_5HT_ expressing mouse geniculate neurons, shown in Figure 1 (Slope of regression line = 0.0000868; Normality test *p* < 0.001; Constant variance test *p* = <0.001; *r*^2^ = 0.0807; *F* = 640.882; *n* = 7,302; *p* < 0.001), Figure 2 (Slope of regression line = 0.0000884; Normality test *p* < 0.001; Constant variance test *p* = <0.001; *r*^2^ = 0.0809; *F* = 228.602; *n* = 2,598; *p* < 0.001), Figure 3 (Slope of regression line = −0.000012516; Normality test *p* < 0.001; Constant variance test *p* = 0.0726; *r*^2^ = 0.0224; *F* = 43.1731; *n* = 1885; *p* < 0.001), and Figure 4 (Slope of regression line = 0.0000356; Normality test *p* < 0.001; Constant variance test *p* < 0.001; *r*^2^ = 0.00968; *F* = 47.128; *n* = 4,823; *p* < 0.001). Plots of ΔF/F_0_ against F_0_ of GRAB_5HT_ expressing human iPSC-derived neuron shown in in Figure 4 (Slope of regression line = 0.00000328; Normality test *p* < 0.001; Constant variance test *p* = <0.001; *r^2^* = 0.000843; *F* = 4.518; *n* = 5,354; *p* = 0.034; Linear regression *t* tests). Plots of ΔF/F_0_ against F_0_ of the GRAB_5HT_ expressing mouse geniculate neurons, shown in Figure S3 for control (Slope of regression line = 0.0000393; Normality test *p* < 0.001; Constant variance test *p* = <0.001; *r*^2^ = 0.0029; *F* = 5.1765; *n* = 1,752; *p* < 0.001), and for TTX (Slope of regression line = 0.00000856; Normality test *p* < 0.001; Constant variance test *p* = <0.001; *r*^2^ = 0.0137; *F* = 23.6859; *n* = 1,705; *p* < 0.001). Plot of ΔF/F_0_ against F_0_ of the GRAB_5HT1.0_ expressing mouse raphe neurons shown in Figure S4 (Slope of regression line = 0.00000980; Normality test *p* < 0.001; Constant variance test *p* < 0.001; *r*^2^ = 0.103; *F* = 81.214; *n* = 711; *p* < 0.001). Plot of ΔF/F_0_ against F_0_ of the GRAB_5HT1.0_ expressing rat hippocampus neurons shown in Figure S5 (Slope of regression line = −0.000033027; Normality test *p* = 0.2222; Constant variance test *p* = 0.0025; *r*^2^ = 0.0426; *F* = 66.3471; *n* = 1,493; *p* < 0.001). (**B**) Fluorescence responses are largely independent of iAChSnFR expression. Plot of ΔF/F_0_ against F_0_ of the iAChSnFR expressing mouse entorhinal astrocyte shown in Figure 5 (Slope of regression line = 0.0000271; Normality test *p* < 0.001; Constant variance test *p* < 0.001; *r*^2^ = 0.00126; *F* = 29.227; n = 23,166; *p* = 0.004). Plot of ΔF/F_0_ against F_0_ of the iAChSnFR expressing mouse MEC neuron shown in Figure S6 (Slope of regression line = 0.0002; Normality test *p* < 0.001; Constant variance test *p* < 0.001; *r*^2^ = 0.0274; *F* = 53.0846; *n* = 7,672; *p* < 0.001).Plot of ΔF/F_0_ against F_0_ of the iAChSnFR expressing mouse pancreatic cell shown in Figure S7 (Slope of regression line = 0.0000228; Normality test *p* < 0.001; Constant variance test *p* < 0.001; *r^2^* = 0.0868; *F* = 549.465; *n* = 5,780; *p* < 0.001.Plot of ΔF/F_0_ against F_0_ of the iAChSnFR expressing mouse adrenal cell shown in Figure S8 (Slope of regression line = −0.0000145; Normality test *p* < 0.001; Constant variance test *p* < 0.001; *r*^2^ = 0.000870; *F* = 6.227; *n* = 7,152; *p* = 0.013; Linear regression *t* tests). (**C**) Fluorescence responses are largely independent of GRAB_NE_, GRAB_OT1.0_, GRAB_DA_, GRAB_His_, OxLight, iGluSnFR, and iGluSnFR3 expression. Plot of ΔF/F_0_ against F_0_ of the GRAB_NE_ expressing mouse amygdalar neuron shown in Figure 6 (Slope of regression line = 0.000437; Normality test *p* = 0.558; Constant variance test *p* = 0.584; *r*^2^ = 0.18064 *F* = 498.670; *n* = 2,279; *p* < 0.001). Plot of ΔF/F_0_ against F_0_ of the GRAB_OT1.0_ expressing mouse paraventricular neuron shown in Figure 7 (Slope of regression line = 0.0000640; Normality test *p* = 0.9767; Constant variance test *p* = 0.0561, *r*^2^ = 0.180; *F* = 237.344; *n* = 1,038; *p* < 0.001) and ventral tegmental neuron shown in Figure 7 (Slope of regression line = −0.0000250; Normality test *p* <0.001; Constant variance test *p* < 0.001; *r*^2^ = 0.0034; *F* = 8.5448; *n* = 2,496; *p* = 0.0035). Plot of ΔF/F_0_ against F_0_ of the GRAB_DA_ expressing mouse amygdalar neuron shown in Figure S9 (Slope of regression line = −0.0000378 (Normality test *p* < 0.001; Constant variance test *p* < 0.001; *r*^2^ = 0.0329; *F* = 177.345; *n* = 5,219; *p* < 0.001). Plot of ΔF/F_0_ against F_0_ of the GRAB_His_ expressing mouse amygdalar neuron shown in Figure S10 (Slope of regression line = −0.0002; Normality test *p* < 0.001; Constant variance test *p* < 0.001; *r*^2^ = 0.0253; *F* = 23.0142; *n* = 886; *p* < 0.001). Plot of ΔF/F_0_ against F_0_ of the OxLight expressing mouse nucleus accumbens neuron shown in Figure S11 (Slope of regression line = −0.0002; Normality test *p* < 0.001; Constant variance test *p* < 0.001; *r*^2^ = 0.0220; *F* = 54.4167; *n* = 2,380; *p* < 0.001). Plot of ΔF/F_0_ against F_0_ of the iGluSnFR expressing mouse amygdalar neuron shown in Figure S12 (Slope of regression line = −0.00000256; Normality test *p* < 0.001; Constant variance test *p* < 0.001; *r*^2^ = 0.0527; *F* = 54.4167; *n* = 10,106; *p* < 0.001). Plot of ΔF/F_0_ against F_0_ of the iGluSnFR3 expressing mouse paraventricular neuron shown in Figure S13 (Slope of regression line = −0.00000103; Normality test *p* < 0.001; Constant variance test *p* < 0.001; *r*^2^ = 0.0271; *F* = 39.0578; *n* = 1,405; *p* < 0.001).

**Table S1.**
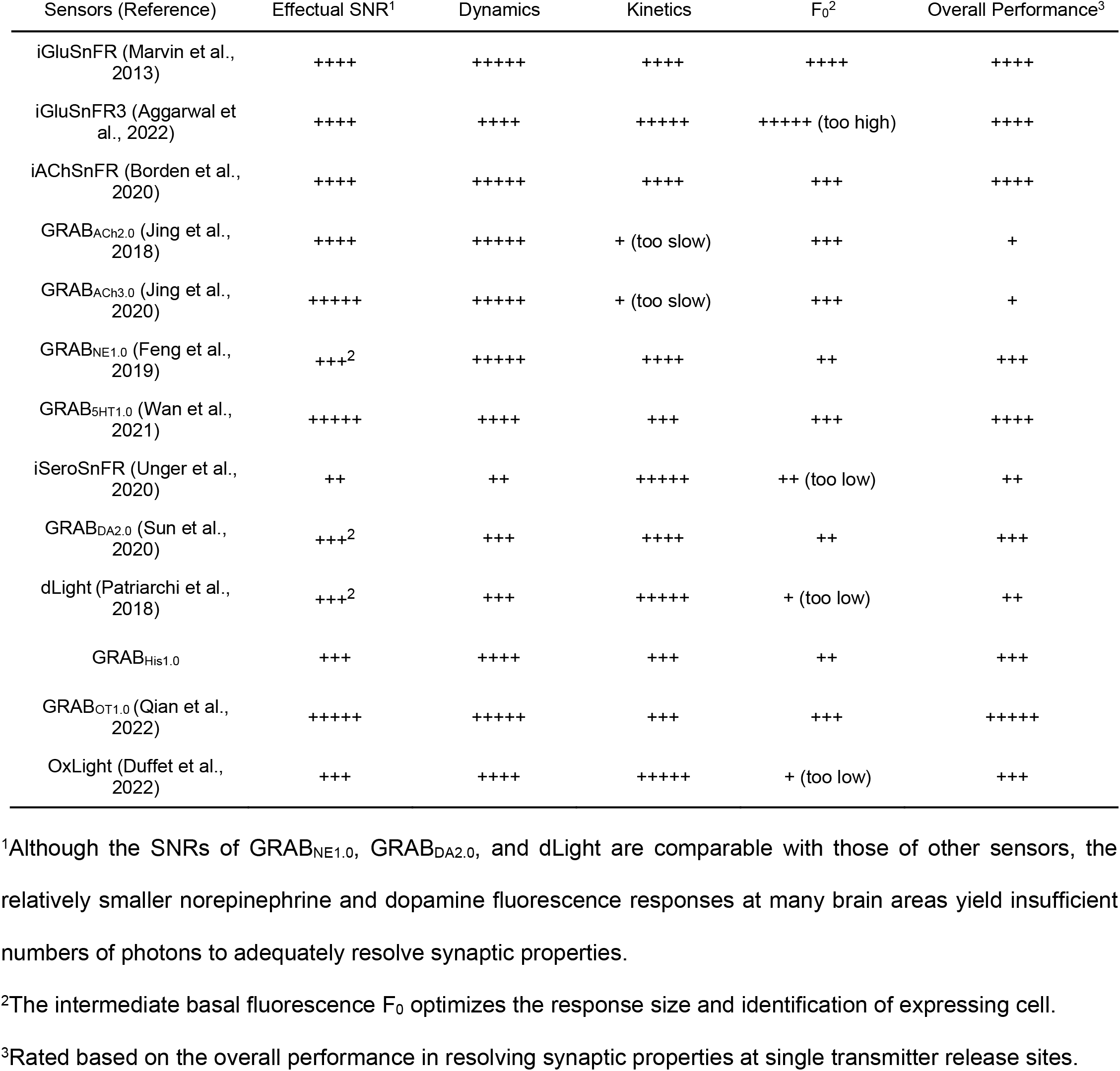
Performance and property rating of genetically encoded sensors.

## Notes

### Competing Interest Statement

The authors have declared no competing interest.

## REFERENCES

Aggarwal A et al. (2022) Glutamate indicators with improved activation kinetics and localization for imaging synaptic transmission. doi: https://www.biorxiv.org/content/10.1101/2022.02.13.480251v1:2022.2002.2013.480251.

Agnati LF, Bjelke B, Fuxe K (1992) Volume transmission in the brain. Am Sci 80:362–373.

Arigovindan M, Fung JC, Elnatan D, Mennella V, Chan YH, Pollard M, Branlund E, Sedat JW, Agard DA (2013) High-resolution restoration of 3D structures from widefield images with extreme low signal-to-noise-ratio. Proceedings of the National Academy of Sciences of the United States of America 110:17344–17349.

Ashford JW (2015) Treatment of Alzheimer’s disease: the legacy of the cholinergic hypothesis, neuroplasticity, and future directions. Journal of Alzheimer’s disease: JAD 47:149–156.

Aston-Jones G, Cohen JD (2005) An integrative theory of locus coeruleus-norepinephrine function: Adaptive gain and optimal performance. Annual review of neuroscience 28:403–450.

Ballinger EC, Ananth M, Talmage DA, Role LW (2016) Basal forebrain cholinergic circuits and signaling in cognition and cognitive decline. Neuron 91:1199–1218.

Barbour B, Hausser M (1997) Intersynaptic diffusion of neurotransmitter. Trends in neurosciences 20:377–384.

Bari A, Xu S, Pignatelli M, Takeuchi D, Feng J, Li Y, Tonegawa S (2020) Differential attentional control mechanisms by two distinct noradrenergic coeruleo-frontal cortical pathways. Proceedings of the National Academy of Sciences of the United States of America.

Bayes A, van de Lagemaat LN, Collins MO, Croning MD, Whittle IR, Choudhary JS, Grant SG (2011) Characterization of the proteome, diseases and evolution of the human postsynaptic density. Nature neuroscience 14:19–21.

Borden PM et al. (2020) A fast genetically encoded fluorescent sensor for faithful in vivo acetylcholine detection in mice, fish, worms and flies. bioRxiv doi: https://doi.org/10.1101/2020.02.07.939504.

Borroto-Escuela DO, Perez De La Mora M, Manger P, Narvaez M, Beggiato S, Crespo-Ramirez M, Navarro G, Wydra K, Diaz-Cabiale Z, Rivera A, Ferraro L, Tanganelli S, Filip M, Franco R, Fuxe K (2018) Brain dopamine transmission in health and Parkinson’s disease: modulation of synaptic transmission and plasticity through volume transmission and dopamine heteroreceptors. Front Synaptic Neurosci 10:20.

Breton-Provencher V, Drummond GT, Feng J, Li Y, Sur M (2022) Spatiotemporal dynamics of noradrenaline during learned behaviour. Nature.

Carter ME, Yizhar O, Chikahisa S, Nguyen H, Adamantidis A, Nishino S, Deisseroth K, de Lecea L (2010) Tuning arousal with optogenetic modulation of locus coeruleus neurons. Nature neuroscience 13:1526–1533.

Dani JA, Bertrand D (2007) Nicotinic acetylcholine receptors and nicotinic cholinergic mechanisms of the central nervous system. Annual review of pharmacology and toxicology 47:699–729.

Darvesh AS, Carroll RT, Geldenhuys WJ, Gudelsky GA, Klein J, Meshul CK, Van der Schyf CJ (2011) In vivo brain microdialysis: advances in neuropsychopharmacology and drug discovery. Expert Opin Drug Dis 6:109127.

Destexhe A, Mainen ZF, Sejnowski TJ (1994) Synthesis of models for excitable membranes, synaptic transmission and neuromodulation using a common kinetic formalism. Journal of computational neuroscience 1:195–230.

Duffet L et al. (2022) A genetically encoded sensor for in vivo imaging of orexin neuropeptides. Nature methods 19:231–241.

Ester M, Kriegel H-P, Sander J, Xu X (1996) A density-based algorithm for discovering clusters in large spatial databases with noise. In: kdd, pp 226–231.

Fatt P, Katz B (1952) Spontaneous subthreshold activity at motor nerve endings. The Journal of physiology 117:109–128.

Feng J, Zhang C, Lischinsky JE, Jing M, Zhou J, Wang H, Zhang Y, Dong A, Wu Z, Wu H, Chen W, Zhang P, Zou J, Hires SA, Zhu JJ, Cui G, Lin D, Du J, Li Y (2019) A genetically encoded fluorescent sensor for rapid and specific in vivo detection of norepinephrine. Neuron 102:745–761 e748.

Foote SL, Bloom FE, Astonjones G (1983) Nucleus locus ceruleus - new evidence of anatomical and physiological specificity. Physiological reviews 63:844–914.

Greengard P (2001) The neurobiology of slow synaptic transmission. Science 294:1024–1030.

Guillaumin MCC, Burdakov D (2021) Neuropeptides as primary mediators of brain circuit connectivity. Frontiers in neuroscience 15:644313.

Henley JM, Wilkinson KA (2016) Synaptic AMPA receptor composition in development, plasticity and disease. Nature reviews Neuroscience 17:337–350.

Herring N, Kalla M, Paterson DJ (2019) The autonomic nervous system and cardiac arrhythmias: current concepts and emerging therapies. Nat Rev Cardiol 16:707–726.

Jackman SL, Regehr WG (2017) The mechanisms and functions of synaptic facilitation. Neuron 94:447–464.

Jensen TP, Zheng K, Cole N, Marvin JS, Looger LL, Rusakov DA (2019) Multiplex imaging relates quantal glutamate release to presynaptic Ca^2+^ homeostasis at multiple synapses in situ. Nature communications 10:1414.

Jing M et al. (2018) A genetically encoded fluorescent acetylcholine indicator for in vitro and in vivo studies. Nature biotechnology 36:726–737.

Jing M et al. (2020) An optimized acetylcholine sensor for monitoring in vivo cholinergic activity. Nature methods 17:1139–1146.

Kaeser PS, Regehr WG (2014) Molecular mechanisms for synchronous, asynchronous, and spontaneous neurotransmitter release. Annual review of physiology 76:333–363.

Kazemipour A, Novak O, Flickinger D, Marvin JS, Abdelfattah AS, King J, Borden PM, Kim JJ, Al-Abdullatif SH, Deal PE, Miller EW, Schreiter ER, Druckmann S, Svoboda K, Looger LL, Podgorski K (2019) Kilohertz framerate two-photon tomography. Nature methods 16:778–786.

Koho S, Tortarolo G, Castello M, Deguchi T, Diaspro A, Vicidomini G (2019) Fourier ring correlation simplifies image restoration in fluorescence microscopy. Nature communications 10:3103.

Labouesse MA, Patriarchi T (2021) A versatile GPCR toolkit to track in vivo neuromodulation: not a one-size-fits-all sensor. Neuropsychopharmacology: official publication of the American College of Neuropsychopharmacology 46:2043–2047.

Landweber L (1951) An iteration formula for fredholm integral equations of the first kind. American Journal of Mathematics 73:615–624.

Li J et al. (2022) A tissue-like neurotransmitter sensor for the brain and gut. Nature 606:94–101.

Lim CS, Kang X, Mirabella V, Zhang H, Bu Q, Araki Y, Hoang ET, Wang S, Shen Y, Choi S, Kaang BK, Chang Q, Pang ZP, Huganir RL, Zhu JJ (2017) BRaf signaling principles unveiled by large-scale human mutation analysis with a rapid lentivirus-based gene replacement method. Genes & development 31:537–552.

Lin L, Gupta S, Zheng WS, Si K, Zhu JJ (2021) Genetically encoded sensors enable micro-and nano-scopic decoding of transmissions in healthy and diseased brains. Molecular psychiatry:443–455.

Marvin JS, Borghuis BG, Tian L, Cichon J, Harnett MT, Akerboom J, Gordus A, Renninger SL, Chen TW, Bargmann CI, Orger MB, Schreiter ER, Demb JB, Gan WB, Hires SA, Looger LL (2013) An optimized fluorescent probe for visualizing glutamate neurotransmission. Nature methods 10:162–170.

Mash DC, Flynn DD, Potter LT (1985) Loss of M2 muscarine receptors in the cerebral cortex in Alzheimer’s disease and experimental cholinergic denervation. Science 228:1115–1117.

Nadim F, Bucher D (2014) Neuromodulation of neurons and synapses. Current opinion in neurobiology 29:48–56.

Neher E (2015) Merits and limitations of vesicle pool models in view of heterogeneous populations of synaptic vesicles. Neuron 87:1131–1142.

Nusbaum MP, Blitz DM, Marder E (2017) Functional consequences of neuropeptide and small-molecule cotransmission. Nature reviews Neuroscience 18:389–403.

Okaty BW, Commons KG, Dymecki SM (2019) Embracing diversity in the 5-HT neuronal system. Nature reviews Neuroscience 20:397–424.

Olive MF, Mehmert KK, Hodge CW (2000) Microdialysis in the mouse nucleus accumbens: a method for detection of monoamine and amino acid neurotransmitters with simultaneous assessment of locomotor activity. Brain Res Protoc 5:16–24.

Patriarchi T, Cho JR, Merten K, Howe MW, Marley A, Xiong WH, Folk RW, Broussard GJ, Liang RQ, Jang MJ, Zhong HN, Dombeck D, von Zastrow M, Nimmerjahn A, Gradinaru V, Williams JT, Tian L (2018) Ultrafast neuronal imaging of dopamine dynamics with designed genetically encoded sensors. Science 360:1420–+.

Picciotto MR, Higley MJ, Mineur yS (2012) Acetylcholine as a neuromodulator: cholinergic signaling shapes nervous system function and behavior. Neuron 76:116–129.

Ponti A, Schwarb P, Gulati A, Baker VJI, Microscopy (2007) Huygens remote manager: a web interface for high-volume batch deconvolution. 9:57–58.

Pulido C, Marty A (2017) Quantal fluctuations in central mammalian synapses: functional role of vesicular docking sites. Physiological reviews 97:1403–1430.

Qian T, Wang H, Wang P, Geng L, Mei L, Osakada T, Tang Y, Kania A, Grinevich V, Stoop R, Lin D, Luo M, Li Y (2022) Compartmental neuropeptide release measured using a new oxytocin sensor. doi: https://www.biorxiv.org/content/10.1101/2022.02.10.480016v1:2022.2002.2010.480016.

Rall W (1967) Distinguishing theoretical synaptic potentials computed for different soma-dendritic distributions of synaptic input. Journal of neurophysiology 30:1138–1168.

Ray S, Naumann R, Burgalossi A, Tang Q, Schmidt H, Brecht M (2014) Grid-layout and theta-modulation of layer 2 pyramidal neurons in medial entorhinal cortex. Science 343:891–896.

Reddy BS, Chatterji BN (1996) An FFT-based technique for translation, rotation, and scale-invariant image registration. IEEE Trans Image Process 5:1266–1271.

Robinson DL, Hermans A, Seipel AT, Wightman RM (2008) Monitoring rapid chemical communication in the brain. Chemical reviews 108:2554–2584.

Sabatini BL, Tian L (2020) Imaging neurotransmitter and neuromodulator dynamics in vivo with genetically encoded indicators. Neuron 108:17–32.

Sage D, Donati L, Soulez F, Fortun D, Schmit G, Seitz A, Guiet R, Vonesch C, Unser M (2017) DeconvolutionLab2: An open-source software for deconvolution microscopy. Methods 115:28–41.

Sarter M, Parikh V, Howe WM (2009) Phasic acetylcholine release and the volume transmission hypothesis: time to move on. Nature reviews Neuroscience 10:383–390.

Satin LS, Kinard TA (1998) Neurotransmitters and their receptors in the islets of Langerhans of the pancreas: what messages do acetylcholine, glutamate, and GABA transmit? Endocrine 8:213–223.

Sauer M (2013) Localization microscopy coming of age: from concepts to biological impact. Journal of cell science 126:3505–3513.

Scammell TE, Arrigoni E, Lipton JO (2017) Neural circuitry of wakefulness and sleep. Neuron 93:747–765.

Schneggenburger R, Meyer AC, Neher E (1999) Released fraction and total size of a pool of immediately available transmitter quanta at a calyx synapse. Neuron 23:399–409.

Small A, Stahlheber S (2014) Fluorophore localization algorithms for super-resolution microscopy. Nature methods 11:267–279.

Sudhof TC (2018) Towards an understanding of synapse formation. Neuron 100:276–293.

Sun FM, Zhou J, Dai B, Qian T, Zeng J, Li X, Zhuo Y, Zhang Y, Wang Y, Qian C, Tan K, Feng J, Dong H, Lin D, Cui G, Li Y (2020) New and improved GRAB fluorescent sensors for monitoring dopaminergic activity in vivo. Nature methods 17.

Sylantyev S, Savtchenko LP, Niu YP, Ivanov AI, Jensen TP, Kullmann DM, Xiao MY, Rusakov DA (2008) Electric fields due to synaptic currents sharpen excitatory transmission. Science 319:1845–1849.

Thomas GJ, Jr., Agard DA (1984) Quantitative analysis of nucleic acids, proteins, and viruses by Raman band deconvolution. Biophys J 46:763–768.

Thompson RE, Larson DR, Webb WW (2002) Precise nanometer localization analysis for individual fluorescent probes. Biophys J 82:2775–2783.

Tomasi C, Manduchi R (1998) Bilateral filtering for gray and color images. In: Sixth international conference on computer vision (IEEE Cat. No. 98CH36271), pp 839–846: IEEE.

Tracy TE et al. (2022) Tau interactome maps synaptic and mitochondrial processes associated with neurodegeneration. Cell 185:712–728 e714.

Ungar A, Phillips JH (1983) Regulation of the adrenal medulla. Physiological reviews 63:787–843.

Unger EK et al. (2020) Directed evolution of a selective and sensitive serotonin biosensor via machine learning. https://papers.ssrn.com/sol3/papers.cfm?abstract_id=3498571 under review in Cell.

Vadodaria KC, Mertens J, Paquola A, Bardy C, Li X, Jappelli R, Fung L, Marchetto MC, Hamm M, Gorris M, Koch P, Gage FH (2016) Generation of functional human serotonergic neurons from fibroblasts. Molecular psychiatry 21:49–61.

Van Der Zee EA, De Jong GI, Strosberg AD, Luiten PG (1993) Muscarinic acetylcholine receptor-expression in astrocytes in the cortex of young and aged rats. Glia 8:42–50.

Vercauteren T, Pennec X, Perchant A, Ayache N (2009) Diffeomorphic demons: efficient non-parametric image registration. Neuroimage 45:S61–72.

Volk L, Chiu SL, Sharma K, Huganir RL (2015) Glutamate synapses in human cognitive disorders. Annual review of neuroscience 38:127–149.

Volkow ND, Michaelides M, Baler R (2019) The neuroscience of drug reward and addiction. Physiological reviews 99:2115–2140.

von Gersdorff H, Borst JGG (2002) Short-term plasticity at the calyx of held. Nature Reviews Neuroscience 3:53–64.

Wan J, Peng W, Li X, Qian T, Song K, Zeng J, Deng F, Hao S, Feng J, Zhang P, Zhang Y, Zou J, Pan S, Shin M, Venton BJ, Zhu JJ, Jing M, Xu M, Li Y (2021) A genetically encoded sensor for measuring serotonin dynamics. Nature neuroscience 24:746–752.

Wang G, Wyskiel DR, Yang W, Wang Y, Milbern LC, Lalanne T, Jiang X, Shen Y, Sun Q-Q, Zhu JJ (2015a) An optogenetics-and imaging-assisted simultaneous multiple patch-clamp recordings system for decoding complex neural circuits. Nature protocols 10:397–412.

Wang G, Bochorishvili G, Chen Y, Salvati KA, Zhang P, Dubel SJ, Perez-Reyes E, Snutch TP, Stornetta RL, Deisseroth K, Erisir A, Todorovic SM, Luo JH, Kapur J, Beenhakker MP, Zhu JJ (2015b) CaV3.2 calcium channels control NMDA receptor-mediated transmission: a new mechanism for absence epilepsy. Genes & development 29:1535–1551.

Weigert M et al. (2018) Content-aware image restoration: pushing the limits of fluorescence microscopy. Nature methods 15:1090–1097.

Wichmann C, Kuner T (2022) Heterogeneity of glutamatergic synapses: cellular mechanisms and network consequences. Physiological reviews 102:269–318.

Wu LG, Hamid E, Shin W, Chiang HC (2014) Exocytosis and endocytosis: modes, functions, and coupling mechanisms. Annual review of physiology 76:301–331.

Wu Z, Lin D, Li Y (2022) Pushing the frontiers: tools for monitoring neurotransmitters and neuromodulators. Nature reviews Neuroscience.

Zemek F, Drtinova L, Nepovimova E, Sepsova V, Korabecny J, Klimes J, Kuca K (2014) Outcomes of Alzheimer’s disease therapy with acetylcholinesterase inhibitors and memantine. Expert opinion on drug safety 13:759–774.

Zhang L, Zhang P, Wang G, Zhang H, Zhang Y, Yu Y, Zhang M, Xiao J, Crespo P, Hell JW, Lin L, Huganir RL, Zhu JJ (2018) Ras and Rap signal bidirectional synaptic plasticity via distinct subcellular microdomains. Neuron 98:783–800.

Zhu PK, Zheng WS, Zhang P, Jing M, Borden PM, Ali F, Guo K, Feng J, Marvin JS, Wang Y, Wan J, Gan L, Kwan AC, Lin L, Looger LL, Li Y, Zhang Y (2020) Nanoscopic visualization of restricted non-volume cholinergic and monoaminergic transmission with genetically encoded sensors. Nano Lett 20:4073–4083.

